# Penicillin Binding Protein Substitutions Co-occur with Fluoroquinolone Resistance in ‘Epidemic’ Lineages of Multi Drug-Resistant *Clostridioides difficile*

**DOI:** 10.1101/2022.05.23.493179

**Authors:** Kate E. Dingle, Jane Freeman, Xavier Didelot, David W. Eyre, Jeremy Swan, William D. Spittal, Emma V. Clark, Keith A. Jolley, A. Sarah Walker, Mark H. Wilcox, Derrick W. Crook

## Abstract

*Clostridioides difficile* remains a key cause of healthcare-associated infection, with multi-drug-resistant (MDR) lineages causing high mortality (≥20%) outbreaks. Cephalosporin treatment is a long-established risk factor, and antimicrobial stewardship a key control. A mechanism underlying raised cephalosporin MICs has not been identified in *C. difficile*, but among other species resistance is often acquired *via* amino acid substitutions in cell wall transpeptidases (penicillin binding proteins, PBPs). Here, we investigated five *C. difficile* transpeptidases (PBP1-5) for recent substitutions. Previously published genome assemblies (n=7096) were obtained, representing sixteen geographically widespread lineages, including healthcare-associated MDR ST1(027), ST3(001) and ST17(018). Recent amino acid substitutions were found within PBP1 (n=50) and PBP3 (n=48), ranging from 1-10 substitutions per genome. β-lactam MICs were measured for closely related pairs of wild-type and PBP substituted isolates separated by 20-273 SNPs. Recombination-corrected, dated phylogenies were constructed to date substitution acquisition. Key substitutions such as PBP3 V497L and PBP1 T674I/N/V emerged independently across multiple lineages. They were associated with extremely high cephalosporin MICs; 1-4 doubling dilutions >wild-type up to ≤1506μg/ml. Substitution patterns varied by lineage and clade, showed geographic structure, and notably occurred post-1990, coincident with the acquisition of *gyrA*/*B* substitutions conferring fluoroquinolone resistance. In conclusion, recent PBP1 and PBP3 substitutions are associated with raised cephalosporin MICs in *C. difficile*. The co-occurrence of resistance to cephalosporins and fluoroquinolones hinders attempts to understand their relative importance in the dissemination of epidemic lineages. Further controlled studies of cephalosporin and fluoroquinolone stewardship are needed to determine their relative effectiveness in outbreak control.

**IMPORTANCE:** Fluoroquinolone and cephalosporin prescribing in healthcare settings have triggered outbreaks of high-mortality, multi-drug resistant *C. difficile* infection. Here, we identify a mechanism of acquired cephalosporin resistance in *C. difficile*, comprising amino acid substitutions in two cell-wall transpeptidase enzymes (penicillin binding proteins). The higher the number of substitutions, the greater the impact on phenotype. Dated phylogenies revealed that resistance to both cephalosporins and fluoroquinolones was co-acquired immediately before clinically important, outbreak strains emerged. PBP substitutions were geographically structured within genetic lineages, suggesting adaptation to local antimicrobial prescribing. Antimicrobial stewardship of cephalosporins and fluoroquinolones is an effective means of *C. difficile* outbreak control. Genetic changes conferring resistance likely impart a ‘fitness-cost’ after antibiotic withdrawal. Our study identifies a mechanism that may explain the contribution of cephalosporin stewardship to resolving outbreak conditions. However, due to the co-occurrence of cephalosporin and fluoroquinolone resistance, further work is needed to determine the relative importance of each.

## INTRODUCTION

*Clostridioides difficile* is among the leading causes of healthcare-associated infection, symptoms ranging from diarrhoea to potentially fatal pseudomembranous colitis (1). Over the last 30 years, unrestricted antimicrobial use has selected multidrug resistant (MDR) *C. difficile* lineages, which can be identified by multilocus sequence type (ST) and/or PCR- ribotype (2-9). Uncontrolled prescribing of antimicrobials such as fluoroquinolones and cephalosporins, which are associated with a high risk of *C. difficile* infection (CDI), creates conditions under which MDR lineages can cause persistent, high-mortality (≥20%) outbreaks (9-18). Such healthcare-associated transmission may be geographically widespread as in the ‘hypervirulent’ ST1(ribotype 027) lineage FQ-R1 (19), and/or prolonged, as in ST17(018), predominating in Japanese and Italian healthcare settings since the 1990s (17, 20). Cases associated with the rapid transmission of MDR lineages are typically superimposed over a background of sporadic, unlinked cases caused by *C. difficile* strains, which lack acquired antimicrobial resistance (21, 22).

Antimicrobial stewardship is an extremely effective means of preventing or resolving CDI outbreaks in healthcare settings (23-29). This approach contributed to the marked decline in fluoroquinolone resistant lineages in the UK a decade ago, with resistant ST1(027), ST3(001), ST42(106) and ST37(017) falling from 67% to ∼3% of cases (30, 31). Fluoroquinolone resistance can be predicted from whole genome sequences by characteristic single nucleotide polymorphisms (SNPs) in the chromosomal *gyrA* and/or *gyrB* genes (30, 32). Equivalent analysis for cephalosporins is lacking, because the genetic mechanism(s) influencing cephalosporin susceptibility in *C. difficile* are unknown. Wild-type MICs are already moderate to high for this species (up to, and >256µg/ml), (33-36), leading to the concept of ‘intrinsic’ rather than acquired cephalosporin resistance (10). For this reason, and the practical difficulties of determining extremely high MICs near the limit of drug solubility, cephalosporin MICs are rarely measured for *C. difficile*.

Studies aiming to understand the mechanism of *C. difficile* cephalosporin resistance have focused on the endogenous *C. difficile* class D β-lactamase, but findings were inconclusive (37, 38). In many bacterial species, reduced susceptibility to cephalosporins and other β-lactams is conferred by amino acid substitutions in penicillin binding proteins (PBPs). These are enzymes catalysing cell wall peptidoglycan biosynthesis, which are classified according to molecular weight (high or low, H/LMW) and enzymatic activity (transpeptidase or carboxypeptidase) (39). β-lactam antibiotics target PBPs by acting as inhibitory substrate analogues (39), binding covalently to the active site serine (40) in the first of three conserved motifs; SXXK, (S/Y)XN and (K/H)(S/T)G (41). β-lactam exposure selects substitutions which reduce the affinity of the drug for the PBP, increasing the MIC (42, 43). However, the nature and frequency of PBP substitutions among clinically important *C. difficile* lineages, and their impact on cephalosporin MIC has not been investigated systematically. Here, our aim was to determine whether PBP substitutions have contributed to the emergence and spread of epidemic MDR *C. difficile*.

## RESULTS

The study was designed as follows. A globally distributed collection of published *C. difficile* genomes was assembled (n=7094) representing sixteen genetic lineages; fourteen commonly associated with CDI in healthcare settings, and two carried asymptomatically (non-toxigenic). The occurrence of recent, within-lineage PBP substitutions was investigated. Cephalosporin (and other β-lactam) MICs were measured for representative strains containing PBP substitutions, and closely related ‘wild-type’ ancestors. Finally, the timing and sequence of PBP substitution and fluoroquinolone resistance acquisition events was investigated phylogenetically.

### Lineages Studied

The 7094 genomes represented lineages ST1(ribotypes 027/198/176/181) (n=1918), ST17(018) (n=279), ST42(106) (n=563), ST3(001) (n=411), ST37(017) (n=424) and its recent descendant ST81(369) (n=39), ST63(053) (n=37), ST54(012) (n=148), ST8(002) (n=593) and its recent descendant ST183 (n=14), ST11(078) (n=628), ST2(014/020) (n=790), ST6(005) (n=404), ST10(015) (n=263), ST7(026) (n=190), ST56(058) (n=16) and two prevalent non-toxigenic genotypes ST26(039) (n=175) and ST15(010) (n=202). Four genetically distinct *C. difficile* clades; 1, 2, 4 and 5 were represented (44). Genomes, accession numbers, AMR predictions from genotype and references are listed per lineage (Table S1).

### C. difficile PBPs

To date, nine PBPs have been described within the *C. difficile* genome (Table 1) five of which are transpeptidases (PBP1-5) (45). HMW PBP1 and PBP3 are essential for growth *in vitro* (46), while LMW PBP2 and PBP4 are not essential for growth, but are required for sporulation (46). Only LMW PBP5 is variably present (45). PBP1-4 were present in all genomes studied, while PBP5 occurred in lineages ST3(001), ST11(078), and ST37(017)/ST81(369). No additional PBPs were identified using known *C. difficile* PBP sequences in low stringency BLAST searches.

**TABLE 1.**
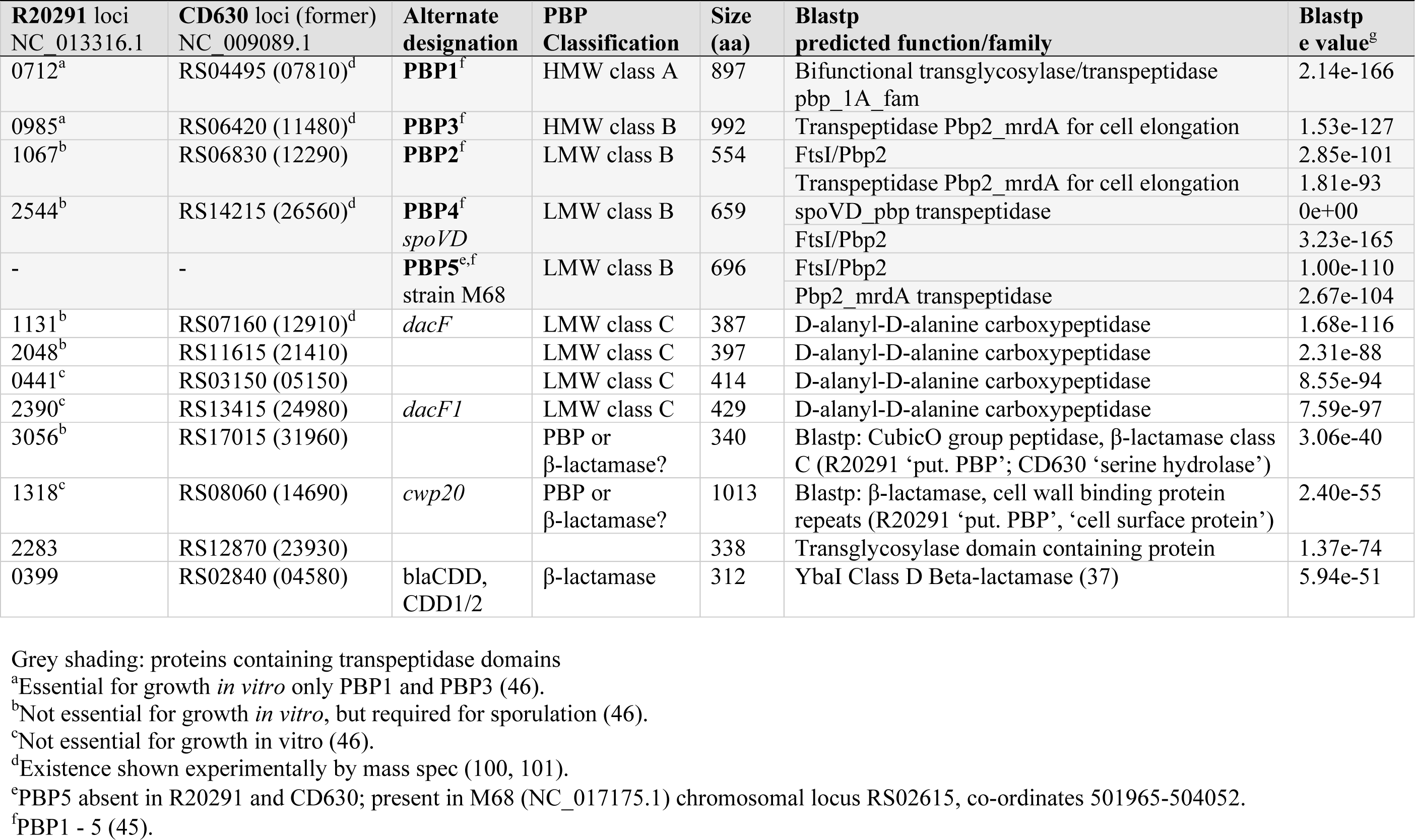
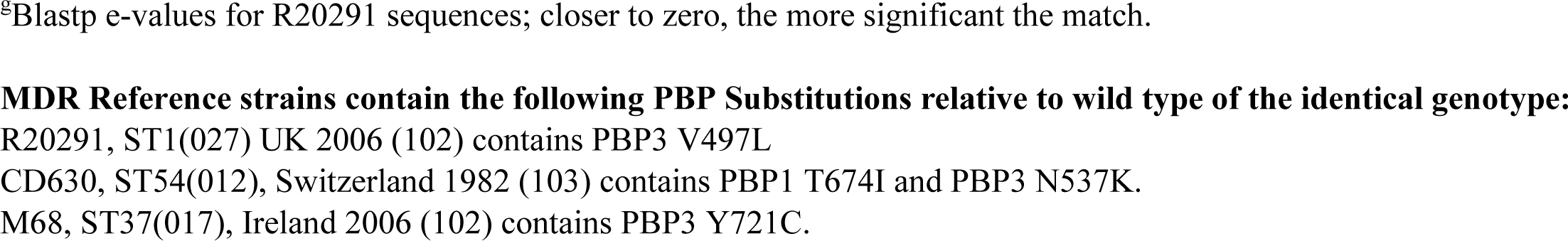
PBPs and β-Lactamases of *C. difficile* Reference Genomes.

| R20291 loci<br>NC 013316.1 | CD630 loci (former)<br>NC 009089.1 | Alternate<br>designation | PBP<br>Classification | Size<br>(aa) | Blastp<br>predicted function/family | Blastp<br>e value<sup>g</sup> |
| --- | --- | --- | --- | --- | --- | --- |
| 0712<sup>a</sup> | RS04495 (07810)<sup>d</sup> | **PBP1**<sup>f</sup> | HMW class A | 897 | Bifunctional transglycosylase/transpeptidase<br>pbp_1A_fam | 2.14e-166 |
| 0985<sup>a</sup> | RS06420 (11480)<sup>d</sup> | **PBP3**<sup>f</sup> | HMW class B | 992 | Transpeptidase Pbp2_mrdA for cell elongation | 1.53e-127 |
| 1067<sup>b</sup> | RS06830 (12290) | **PBP2**<sup>f</sup> | LMW class B | 554 | FtsI/Pbp2 | 2.85e-101 |
|  |  |  |  |  | Transpeptidase Pbp2_mrdA for cell elongation | 1.81e-93 |
| 2544<sup>b</sup> | RS14215 (26560)<sup>d</sup> | **PBP4**<sup>f</sup><br>*spoVD* | LMW class B | 659 | spoVD_pbp transpeptidase | 0e+00 |
|  |  |  |  |  | FtsI/Pbp2 | 3.23e-165 |
| - | - | **PBP5**<sup>e,f</sup><br>strain M68 | LMW class B | 696 | FtsI/Pbp2 | 1.00e-110 |
|  |  |  |  |  | Pbp2_mrdA transpeptidase | 2.67e-104 |
| 1131<sup>b</sup> | RS07160 (12910)<sup>d</sup> | *dacF* | LMW class C | 387 | D-alanyl-D-alanine carboxypeptidase | 1.68e-116 |
| 2048<sup>b</sup> | RS11615 (21410) |  | LMW class C | 397 | D-alanyl-D-alanine carboxypeptidase | 2.31e-88 |
| 0441<sup>c</sup> | RS03150 (05150) |  | LMW class C | 414 | D-alanyl-D-alanine carboxypeptidase | 8.55e-94 |
| 2390<sup>c</sup> | RS13415 (24980) | *dacF1* | LMW class C | 429 | D-alanyl-D-alanine carboxypeptidase | 7.59e-97 |
| 3056<sup>b</sup> | RS17015 (31960) | | PBP or<br>β-lactamase? | 340 | Blastp: CubicO group peptidase, β-lactamase class<br>C (R20291 'put. PBP'; CD630 'serine hydrolase') | 3.06e-40 |
| 1318<sup>c</sup> | RS08060 (14690) | *cwp20* | PBP or<br>β-lactamase? | 1013 | Blastp: β-lactamase, cell wall binding protein<br>repeats (R20291 'put. PBP', 'cell surface protein') | 2.40e-55 |
| 2283 | RS12870 (23930) |  |  | 338 | Transglycosylase domain containing protein | 1.37e-74 |
| 0399 | RS02840 (04580) | blaCDD,<br>CDD1/2 | β-lactamase | 312 | YbaI Class D Beta-lactamase (37) | 5.94e-51 |

Grey shading: proteins containing transpeptidase domains
<sup>a</sup>Essential for growth *in vitro* only PBP1 and PBP3 (46).
<sup>b</sup>Not essential for growth *in vitro*, but required for sporulation (46).
<sup>c</sup>Not essential for growth *in vitro* (46).
<sup>d</sup>Existence shown experimentally by mass spec (100, 101).
<sup>e</sup>PBP5 absent in R20291 and CD630; present in M68 (NC\_017175.1) chromosomal locus RS02615, co-ordinates 501965-504052.
<sup>f</sup>PBP1 - 5 (45).
<sup>g</sup>Blastp e-values for R20291 sequences; closer to zero, the more significant the match.
**MDR Reference strains contain the following PBP Substitutions relative to wild type of the identical genotype:**

### Recent PBP Substitutions

PBP gene sequences were compared within each lineage to identify recent single nucleotide polymorphisms (SNPs). These were absent or rare in LMW PBP2, PBP4 and PBP5. In contrast, multiple SNPs occurred in HMW PBP1 and PBP3, almost all of which were non-synonymous. The resultant amino acid substitutions affected a total of 48/993 (4.8%) positions in PBP3 and 50/856-926 (5.8-5.4%) positions in PBP1 (variable PBP1 size due to C-terminal repeats). Substitution data are shown per isolate (Table S1).

The frequency of each amino acid substitution was recorded per lineage (Table 2). This identified the most common substitutions within and among lineages. In PBP3, V497L was most frequent, occurring in 2897 genomes and 10 lineages, followed by A778V in 664 genomes and 10 lineages. In PBP1, T674I/N/V was most frequent, occurring in 1379 genomes of 10 lineages, followed by A555T in 442 genomes of 7 lineages. These data were then plotted to visualise the relative positions and frequencies of substitutions within PBP1 and PBP3 (Figure 1). Virtually all substitutions occurred within the conserved transpeptidase domains, flanking the active site motifs.

**Figure 1.**
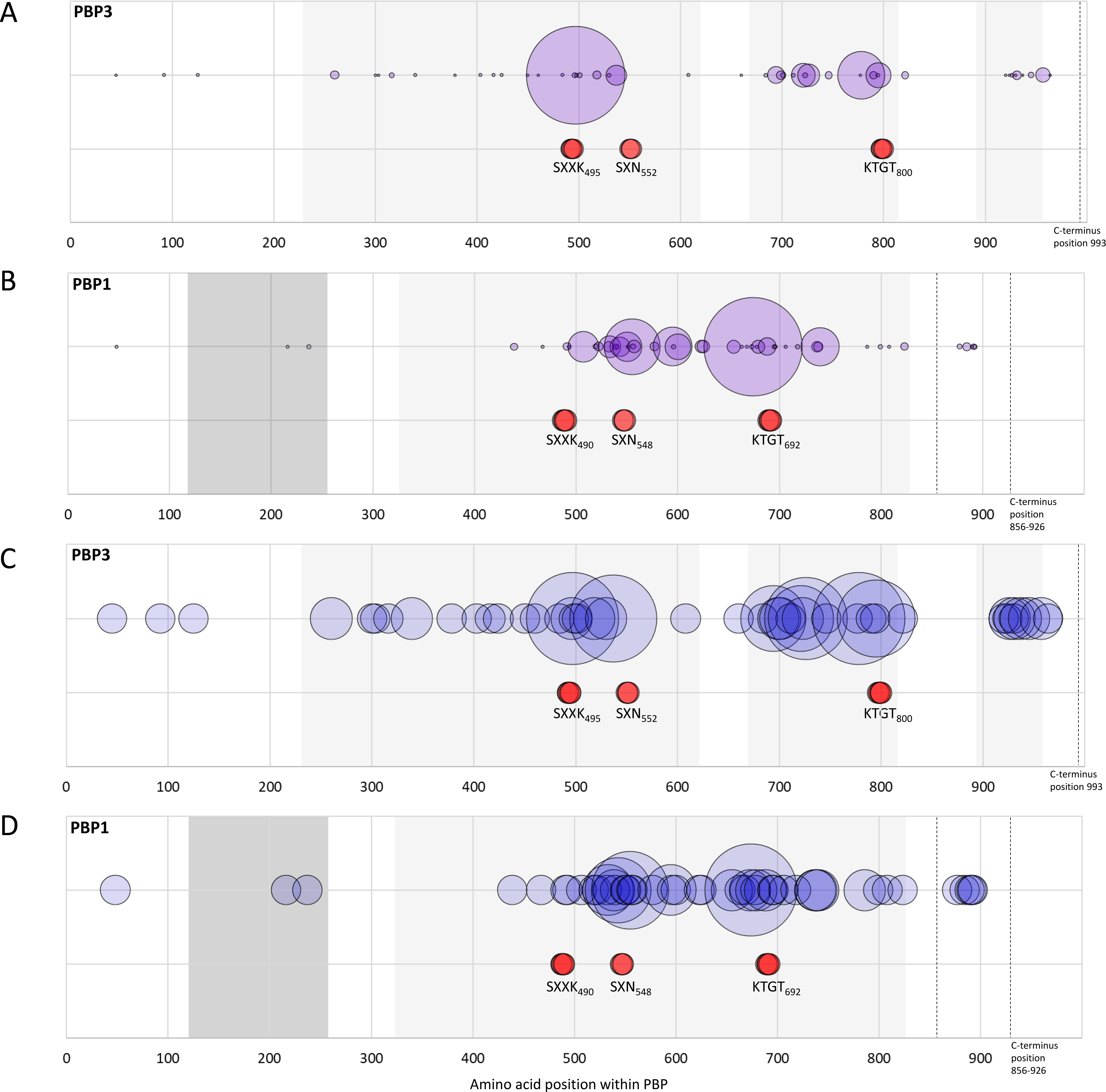
Positions and relative frequency of amino acid substitutions within PBP3 (993 amino acids long) and PBP1 (856-926 amino acids). (A) Substitutions within PBP3 (n=48) are represented by purple circles, plotted according to locations within the PBP (x-axis). Circles are scaled according to substitution frequency within the entire dataset. Light grey shading indicates the position of conserved transpeptidase domains identified by BLASTP. Red circles indicate transpeptidase catalytic motifs. (B) As (A), but PBP1 (n substitutions = 50), and dark grey indicates N-terminal glycosyl transferase domain. (C) As (A), except relative sizes of blue circle indicates the number of lineages in which each substitution was identified. (D) As (B), blue circles again indicating the number of lineages in which each substitution was identified. Raw data, including the identity of each substitution are shown in Table 2.

**Table 2.**
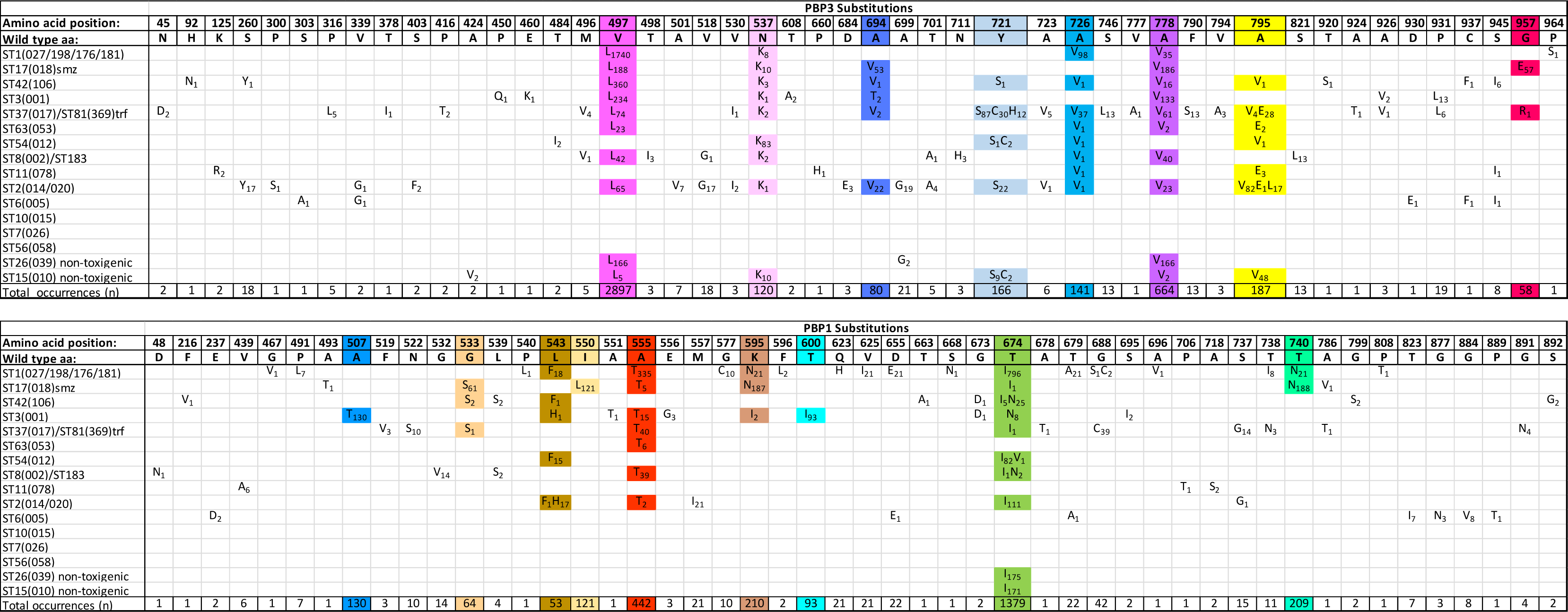
PBP3 and PBP1 substitutions identified within each of the sixteen lineages studied. Upper panel PBP3, lower panel PBP1, with lineages in the left hand column. ‘Amino acid position’ across the top of each panel indicates position within PBP3 or PBP1, together with the wild type amino acid. Amino acid substitutions are indicated within each panel; their frequency within-lineage shown in subscript. The overall frequency of each substitution within the entire data set is indicated in the bottom row. Coloured boxes indicate the substitutions occurring over 50 times within the dataset.

### Co-occurrence of PBP substitutions with fluoroquinolone resistance

In addition to PBP substitutions, the presence of fluoroquinolone resistance was investigated in all sixteen lineages (Figure 2, Table S1). Their co-occurrence was striking in lineages ST1(027/198/176/181), ST17(018), ST42(106), ST3(001), ST37(017)/ST81(369), ST63(053), ST54(012), ST8(002)/ST183 (Figure 2), suggesting the possibility of almost simultaneous acquisition. The presence of further AMR determinants, *ermB* (clindamycin resistance) and, or *rpoB* substitutions (rifampicin resistance), was also recorded for each genome (Table S1).

**Figure 2.**
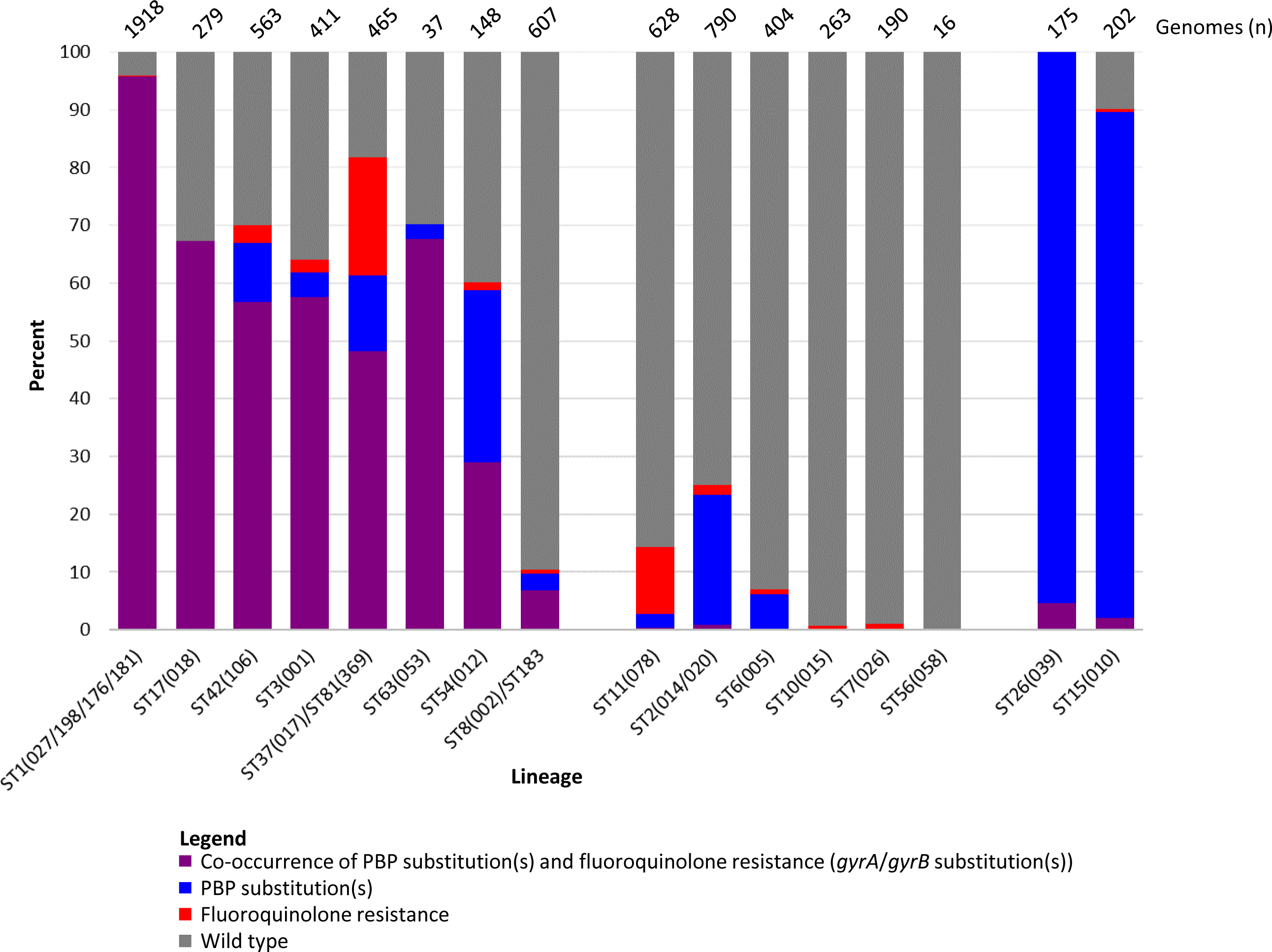
Occurrence of PBP substitutions and fluoroquinolone resistance in the 16 lineages studied.

Among the fourteen clinically important lineages, PBP substitutions occurred relatively rarely in the absence fluoroquinolone resistance (Figure 2). In this respect ST2(014/020) (178/790 genomes, 22.5%), ST54(012) (46/148 genomes, 31.1%) and recent ST42(106) USA genomes (44/180, 24.4%) were noteworthy. Interestingly, PBP substitutions occurred without fluoroquinolone resistance in the majority of genomes belonging to the non-toxigenic lineages ST26(039) (167/175, 95.4%) and ST15(010) (182/202 (90.1%), (Figure 2, Table S1).

### Impact of PBP substitutions on β-lactam MIC

β-lactam MICs were measured for isolates representing eight of the clinically important lineages studied. Four of these lineages contained both ‘wild-type’ and PBP-substituted strains; ST1(027), (both FQ-R1 and FQ-R2 (19)), ST17(018), ST3(001) and ST42(106). The remaining four lineages, included for comparison, were ‘wild-type’ only, lacking PBP substitutions; ST10(015), ST6(005), ST56(058) and ST7(026).

The choice of isolates from ST1(027), ST17(018), ST3(001) and ST42(106), was based on low numbers of SNP differences between wild type and PBP substituted genomes (Figure 3A). In total, ten different PBP substitutions were represented, three in PBP3 and seven in PBP1 (Figure 3B), including the four most frequently identified substitutions. Isolates containing PBP substitutions showed increased cephalosporin and carbapenem MICs relative to wild-type, but their penicillin MICs were unchanged (Figure 3A). The greatest increases in cephalosporin MICs were associated with the highest numbers of substitutions, for example cefuroxime MIC increased from 376 to 1506μg/ml in ST3(001) (four substitutions) and ST17(018) (five substitutions). Cephradine MIC increased from 36 to 239μg/ml in the latter. Intriguingly, the wild-type ancestors of these four PBP substituted lineages had cefotaxime MICs which were still higher than the four lineages which have not yielded PBP substituted strains; cefuroxime 128µg/ml vs 376µg/ml and cefotaxime 128µg/ml vs 256µg/ml.

**Figure 3.**
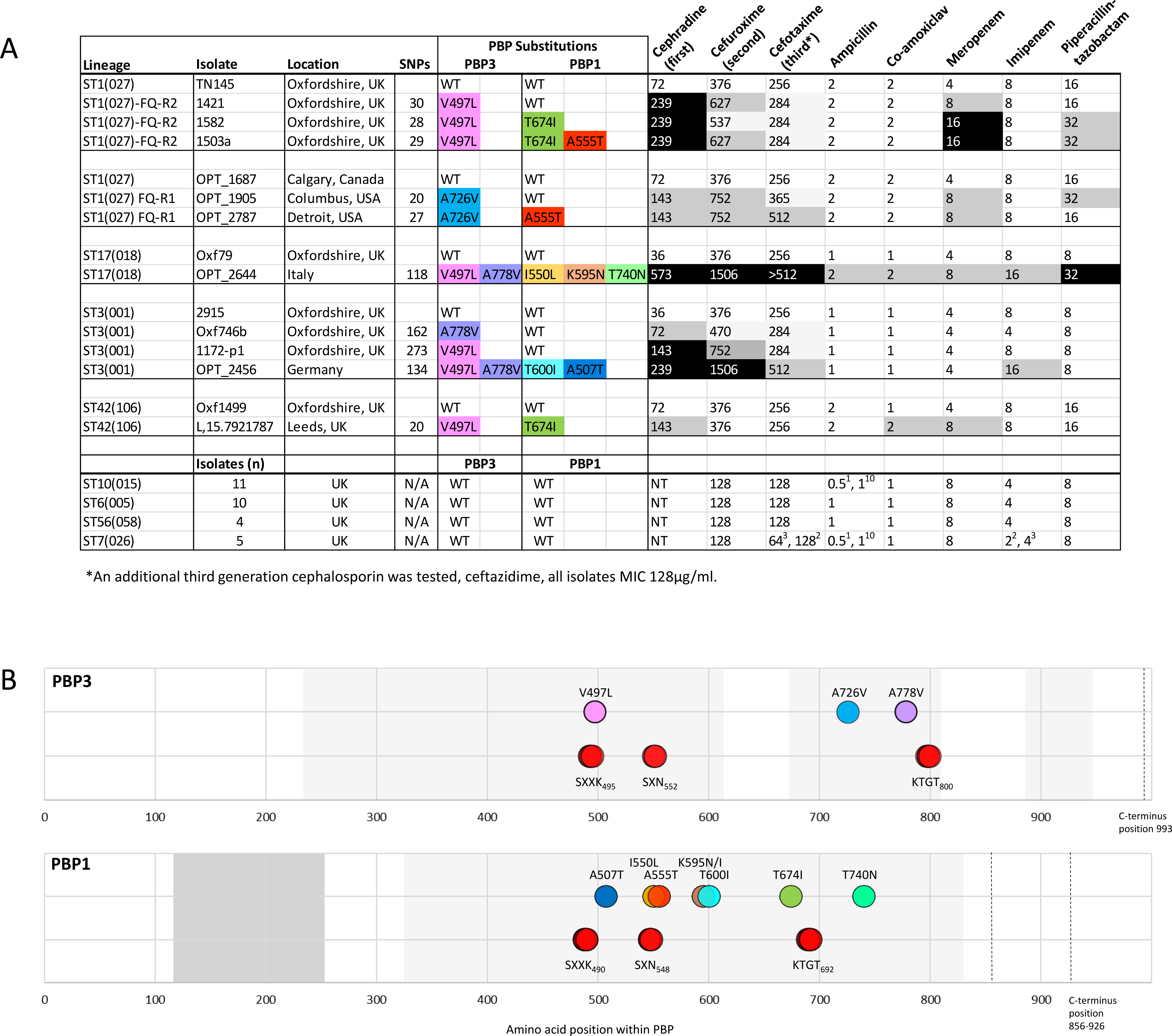
β-lactam MICs of wild type and PBP substituted *C. difficile* isolates. (A) MICs measured for the lineages, strains and β-lactams shown. PBP substitutions are highlighted by colour. Intensity of grey shading indicates fold increase in MIC from wild type. Numbers in superscript indicate the numbers of isolates tested, if >1. WT: wild-type, NT: not tested. (B) Positions of the PBP substitutions (coloured as in (A)) contained in the isolates that underwent phenotyping (A), relative to the conserved transpeptidase domains (grey) and the active site motifs (red circles).

### Phylogenetic analyses

Recombination-corrected phylogenies were constructed to date and identify the sequence in which PBP substitution and fluoroquinolone resistance were acquired by seven genetic lineages. These included the four PBP substituted lineages which had been phenotyped (ST1(027), ST17(018), ST3(001) and ST42 (106)), and a further three lineages containing notable MDR PBP substituted fluoroquinolone resistant strains. These were ST8(002)/ST183 and ST37(017)/ST81 - both important in South East Asia, and ST54(012) - notable in Costa Rica (47-51) (Figures 4-7).

**Figure 4.**
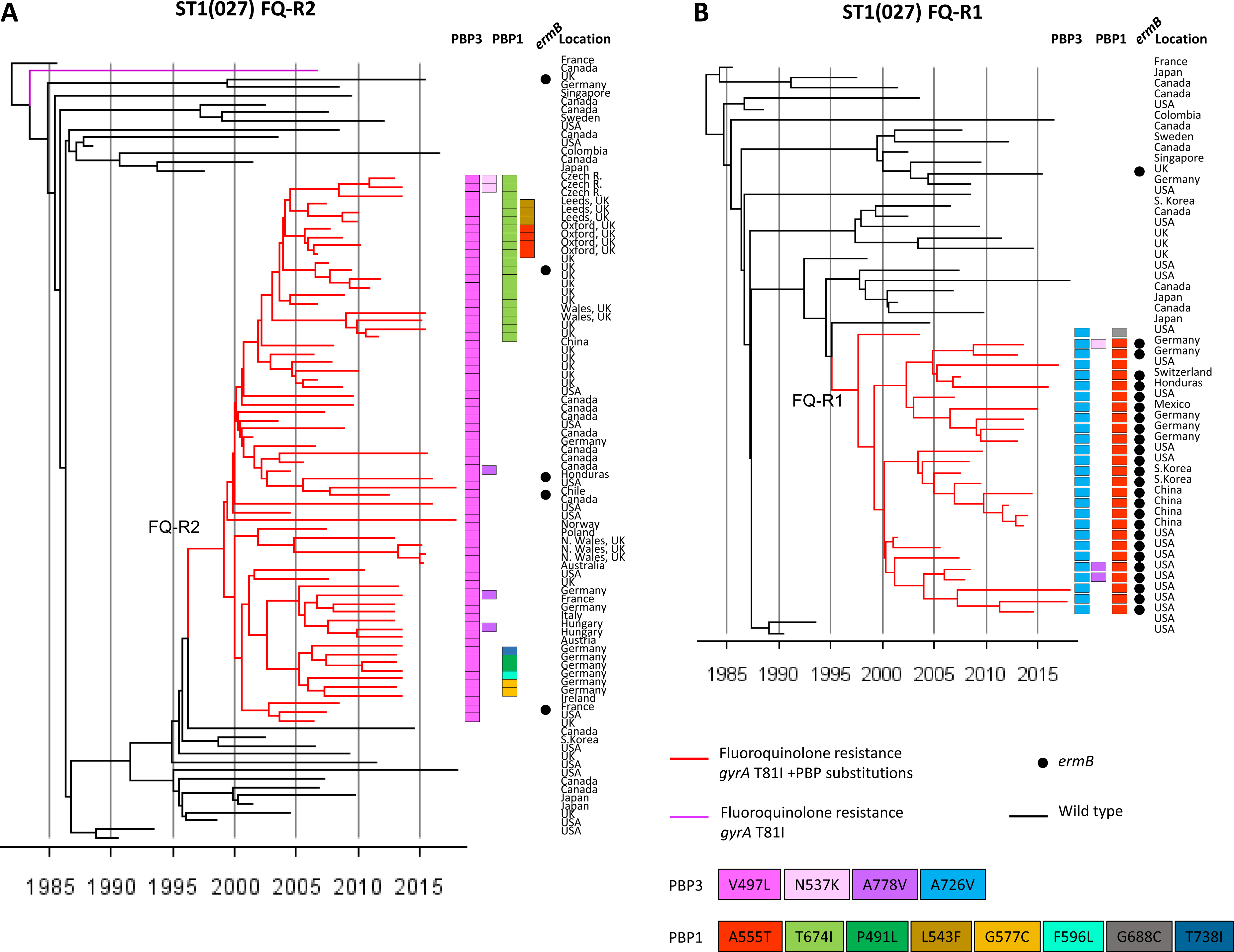
Phylogenetic analysis of lineage ST1(027) (A) Dated phylogeny to show the emergence of lineage ST1(027) FQ-R1 (n=27) (19) (red branches) from wild type (n=27). (B) Dated phylogeny to show the emergence of lineage ST1(027) FQ-R2 (n=67) (19) (red branches) from wild type (n=27). Genomes were chosen to maximise temporal and geographic spread of wild type and AMR strains, and to represent the diversity of PBP substitutions detected. AMR determinants and PBP substitutions are as indicated in the key. Co-occurrence of fluoroquinolone resistance and PBP substitutions is highlighted by red branches. PBP substitutions are highlighted by coloured squares.

Irrespective of lineage, the sequence of PBP substitution acquisitions in epidemic strains typically started with PBP3 V497L and/or A778V (ie. the most frequent PBP3 substitutions, Table 2). Then further PBP substitutions followed, yielding a variety of patterns. Among lineages well known for epidemic spread, the initial PBP3 V497L substitution occurred simultaneously with fluoroquinolone resistance (Figures 4-7). One notable exception was the ST1(027) FQ-R1 lineage in which PBP3 A726V substitution occurred first, while PBP3 V497L was absent (Figure 4B).

PBP substituted, fluoroquinolone resistant clades evolved more than once in ST1(027), ST3(001), ST37(017)/ST81(369), ST42(106), and ST54(012) (Figures 4A,B, 5B, 6B, 7A,B). Their PBP substitution patterns each showed geographic structure (Figures 4-8). MDR clades with the highest numbers of PBP substitutions were identified within ST17(018) in Italy and South East Asia (Figure 5A), ST3(001) in UK/Germany (Figure 5B), ST8(002)/ST183 in Japan (Figure 6A), and ST37(017)/ST81(369) in South East Asia (Figure 6B).

**Figure 5.**
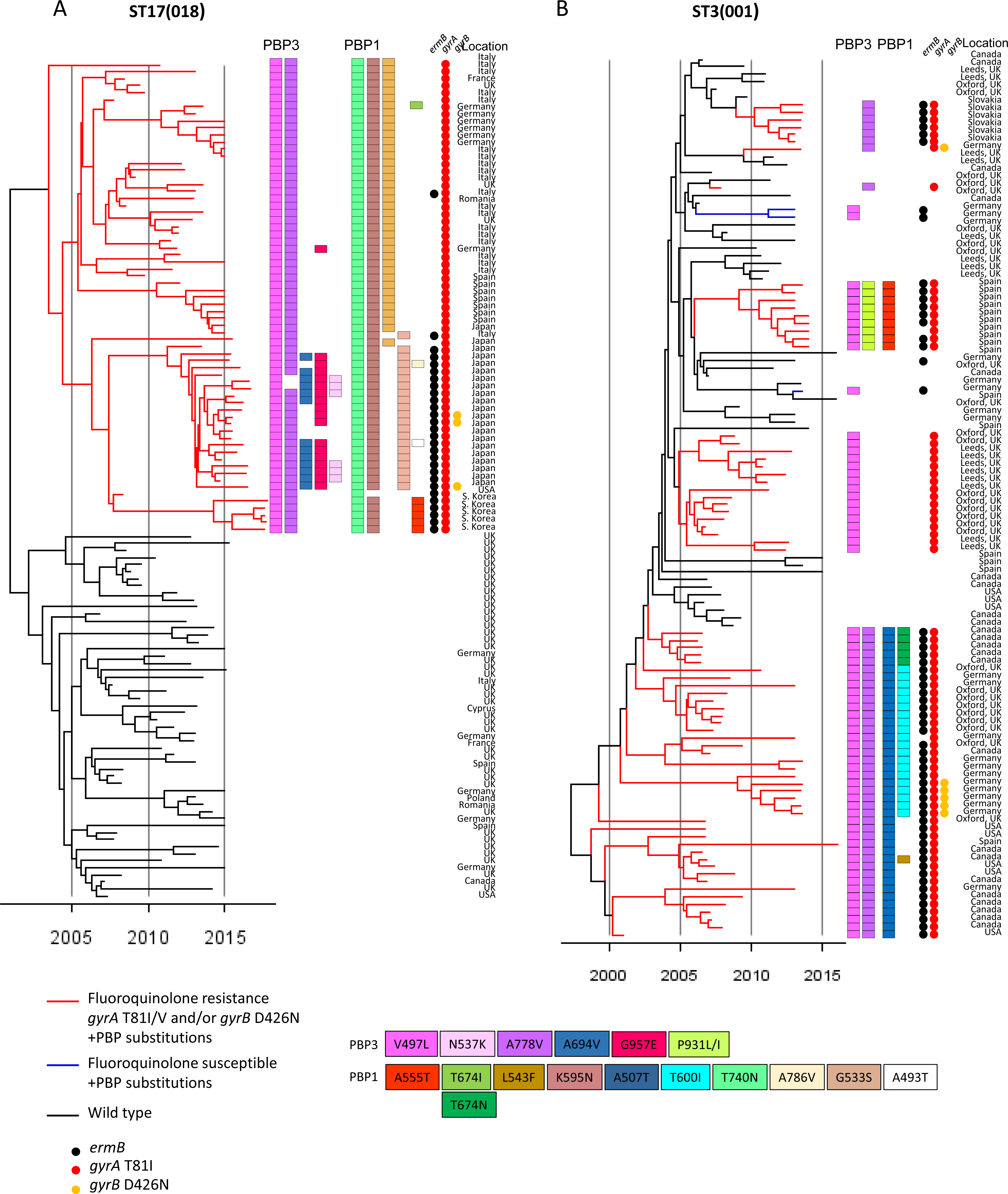
Phylogenetic analysis of lineages ST17(018) and ST3(001) (A) Dated phylogeny to show the emergence of MDR lineage ST17(018) (n=66) (red branches) from wild type (n=53). MDR strains from Europe, South East Asia and North America were chosen to maximise geographic spread and PBP substitutions. These and other AMR determinants and PBP substitutions are as indicated in the key. Co-occurrence of fluoroquinolone resistance and PBP substitutions is highlighted by red branches. (B) Dated phylogeny showing the evolutionary relationship of wild type (n=40) and PBP substituted ST3(001) genomes (n=77).

**Figure 6.**
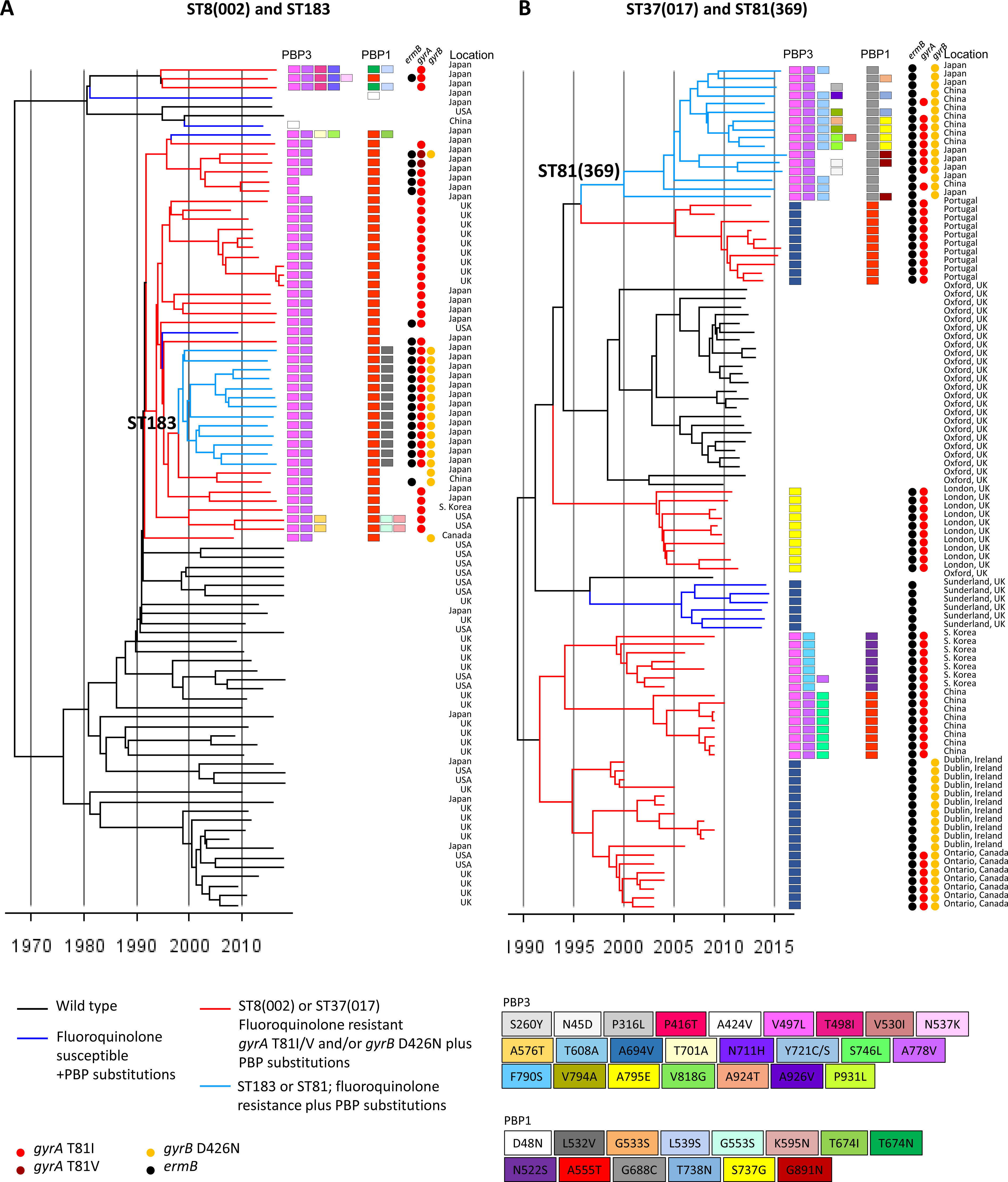
Phylogenetic analysis of lineages ST37(017)/ST81(369) and ST8(002)/ST183. (A) Dated phylogeny showing the evolutionary relationship of wild type (n=43) and PBP substituted ST8(002)/ST183 strains (n=47). Wild type genomes were chosen to maximise genetic diversity (inferred using a previously constructed phylogeny (Dingle et al., 2017)). The MDR ST183 clade was identified here as emergent from MDR ST8(002). Co-occurrence of fluoroquinolone resistance and PBP substitutions is indicated by red branches in ST8(002) and light blue branches in ST183. (B) Dated phylogeny showing the evolutionary relationship of wild type (n=25) and PBP substituted ST37(017) (n=59) and ST81(369) (n=16) genomes. The MDR ST37(017) genomes were chosen to include representatives of strains associated with well documented outbreaks and other potential clusters identified by location and PBP substitution patterns. The MDR ST81(369) clade was identified here as emergent from MDR ST37(017). The majority of available wild type ST37(017) genomes represented healthcare-associated and asymptomatically carried (infant) strains from a single location (Oxfordshire, UK (30; 104)). Co-occurrence of fluoroquinolone resistance and PBP substitutions is indicated by red branches in ST37(017) and light blue branches in ST81.

**Figure 7.**
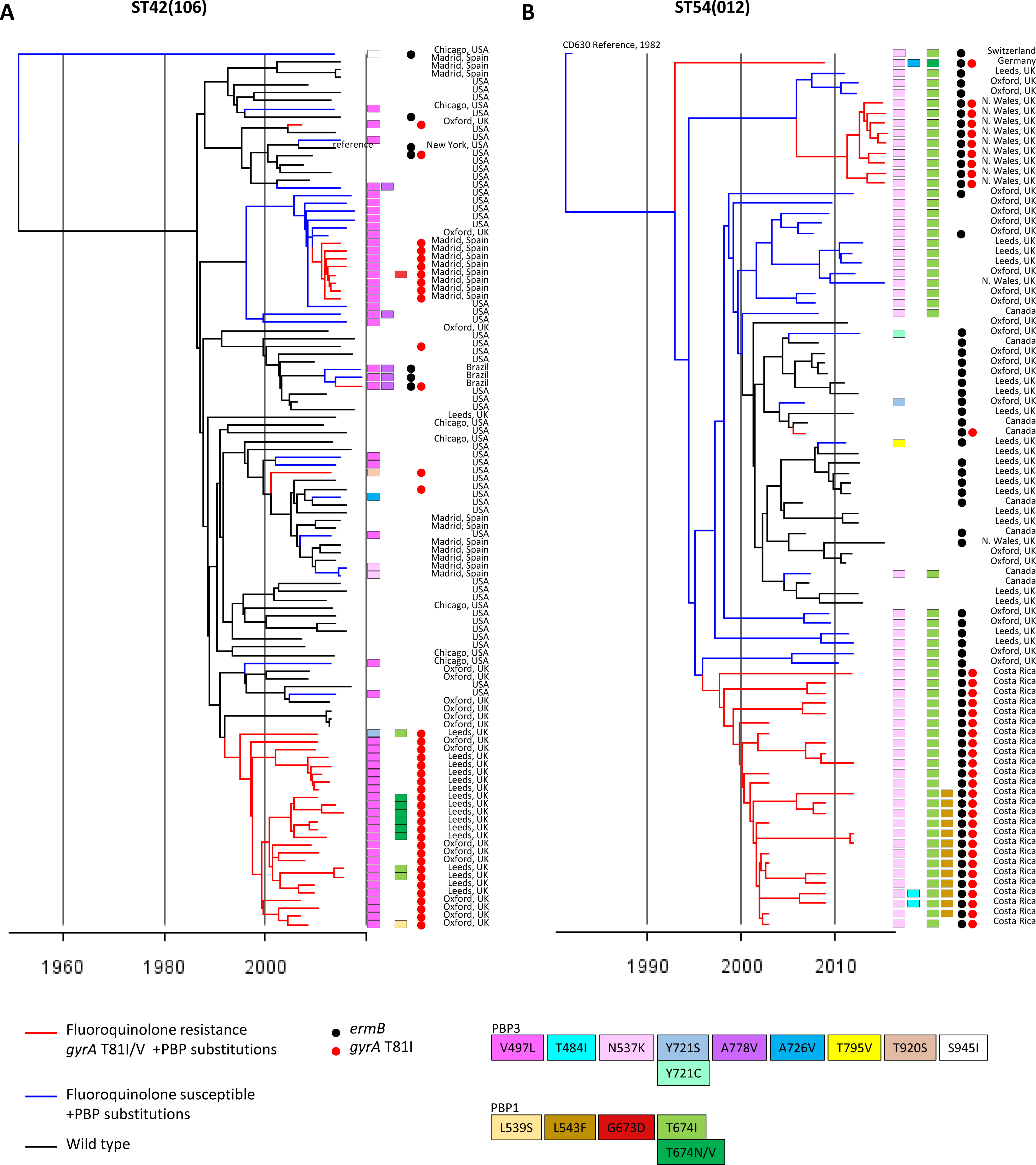
Phylogenetic analysis of lineages ST42(106) and ST54(012) (A) Dated phylogeny for the ST42(106) lineage (n=111 genomes) showing the evolutionary relationship of wild type and fluoroquinolone and/or PBP substituted strains, chosen to represent overall diversity in terms of locations and PBP substitution patterns. (B) As above, but for the ST54(012) lineage (n=107 genomes). In both (A) and (B) the co-occurrence of fluoroquinolone resistance and PBP substitutions indicated by red branches, and PBP substitutions alone by blue branches.

**Figure 8.**
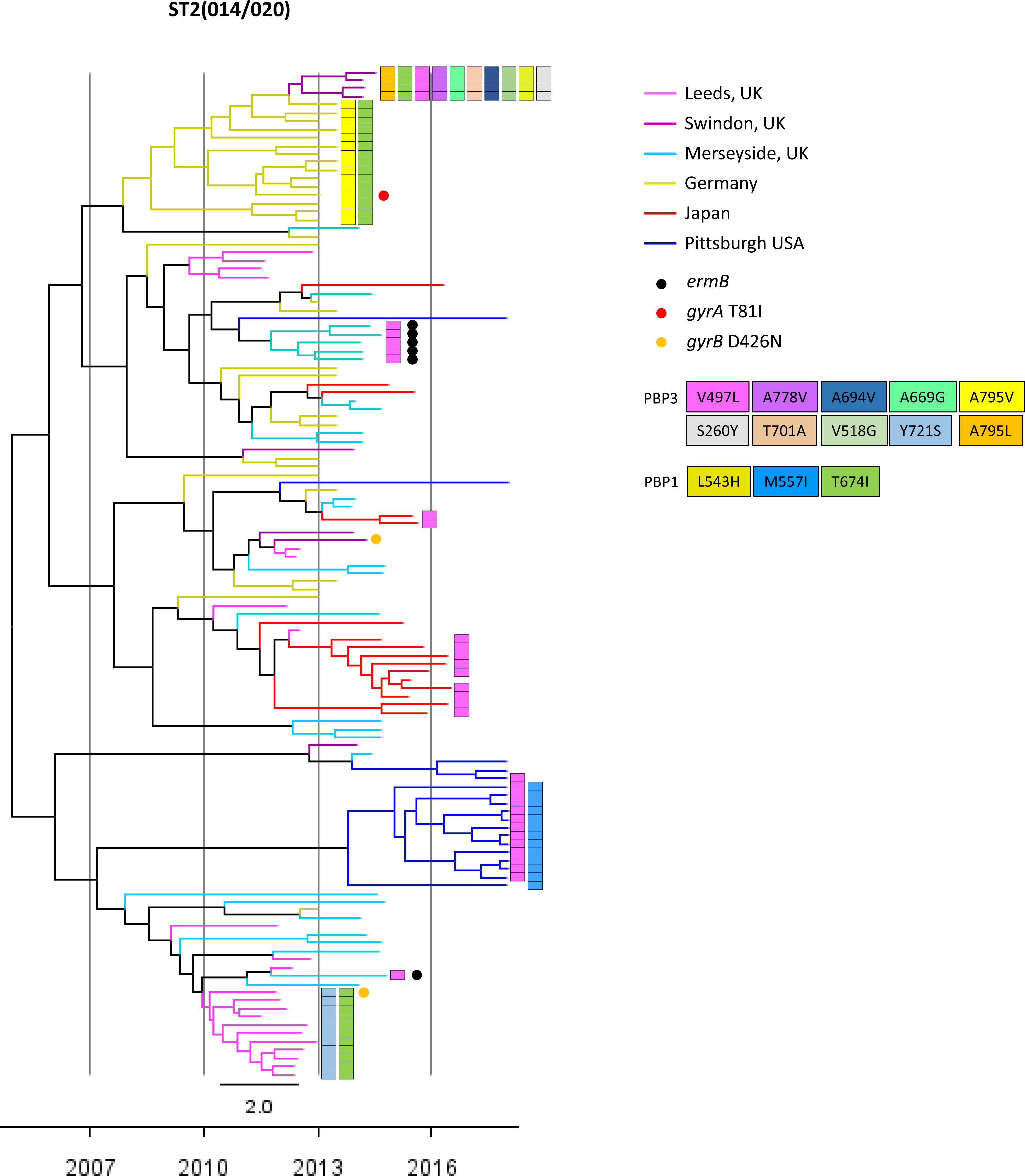
Phylogenetic analysis of ST2(014/020) genomes. Dated phylogeny showing the evolutionary relationship between wild type (n=63) and PBP substituted ST2(014/020) strains (n=60) lacking fluoroquinolone resistance. Branch colour indicates one of six locations. Clustering of PBP substituted genomes is compared with wild type for six independent geographic locations. Wild type genomes from each location were collected concurrently with the PBP substituted strains. Occurrence of AMR determinants is indicated as shown in the key.

The occurrence of PBP substitutions in the absence of fluoroquinolone resistance was investigated phylogenetically in ST2(014/020) (Figure 8), and to a lesser extent in ST42(106) and ST54(012) (Figure 7). For ST2(014/020), a dated phylogeny was constructed using PBP substituted genomes from six independent locations, together with wild type genomes from the same locations and dates (Figure 8). The PBP substituted genomes clustered by location, while the wild type strains did not. Interestingly, both ST42(106) and ST54(012) phylogenies (Figure 7) contained clades where PBP substitution acquisition preceded fluoroquinolone resistance. ST42(106) recently replaced ST1(027) as the most prevalent lineage in North America (Carlson et al., 2020), but a single MDR clade was not apparent; PBP3 V497L occurred on multiple independent occasions within the ST42(106) phylogeny.

### Evolutionary mechanisms of PBP substitution acquisition

The almost total absence of non-synonymous SNPs within each lineage suggested that PBP substitutions accumulate by the step-wise fixation of *de novo* point mutations in response to β-lactam selection, rather than the import of novel variants by horizontal genetic exchange. However, two important recombination events were identified by the co-import of synonymous and non-synonymous SNPs, which changed PBP variants within toxin A-B+ ST81(369) (Figure S1) and in ST1(181), relative to their ancestral wild type A-B+ ST37(017) and ST1(027) wild type respectively. The ST81(369) PBP3 gene was acquired as part of a very long (∼150kb) recombination event, the donor being closely related to ST8(002) (Figure S1). In ST1(181), described in Greece and Romania (52, 53) recombination events affected both PBP1 and PBP3, but their ∼400kb separation around the chromosome suggests the events were independent; (i) the PBP3 allele (100) was identical to clade 1 ST17(018), indicating inter clade 1/2 recombination, and (ii) the PBP1 allele (360) contained 14 SNPs and 6 amino acid differences relative to wild type ST1.

## DISCUSSION

To date, studies aiming to determine β-lactam resistance mechanisms in *C. difficile* have focused on the endogenous *C. difficile* class D β-lactamase (37, 38). PBP substitutions have been reported only occasionally, associated with raised carbapenem MICs in a single lineage (45, 54). PBPs with reduced β-lactam affinity are clinically important in other Gram-positive pathogens, for example *S. pneumoniae* (43), and methicillin resistant *Staphylococcus aureus* (55). The present study was therefore performed to investigate systematically, and phenotypically, recent PBP substitutions among clinically important lineages of *C. difficile*.

We identified multiple, recent PBP substitutions, which are focused in the conserved functional domains of the two HMW *C. difficile* transpeptidases, PBP1 and PBP3 (Table 2, Figure 1). Substitutions were associated with raised cephalosporin MICs, relative to closely related wild type strains, and the higher the number of substitutions, the higher the MIC (Figure 3A). The mechanism underlying substitution acquisition was not recombination, but rather the accumulation of *de novo* chromosomal mutations in response to selective pressure. This was indicated because virtually all SNPs were non-synonymous, flanked the catalytic domains (Figure 1), and arose multiple times (Table 2). Only two major recombination events were found involving ST81(369) (Figure S1), and ST1(181). Although PBP5 transpeptidase is variably present, it was not recently acquired by MDR *C. difficile* lineages. Its constant chromosomal location suggests that gradual loss, rather than recent acquisition may explain its variable presence.

The co-occurrence of PBP substitutions with fluoroquinolone resistance in the clinically important epidemic lineages was striking (Figure 2). This suggests that cephalosporin stewardship may be equal to fluoroquinolone stewardship (30) in its effectiveness for outbreak control. Furthermore, simultaneous stewardship of both drugs may have a greater impact on the control of MDR lineages, than either drug alone. Studies performed over 20 years ago, before widespread fluoroquinolone resistance emerged in *C. difficile*, reported cephalosporin stewardship alone to be successful (24, 56, 57). Approximately 20 years elapsed between the introduction of first generation cephalosporins (mid 1960s) and fluoroquinolones (late 1980s) (58). Selection of the wild-type precursors of the current epidemic lineages (Figure 3A), with cephalosporin MICs that exceed the wild type of other lineages, (WT cefuroxime 128µg/ml vs 376µg/ml and WT cefotaxime 128µg/ml vs 256µg/ml) may have occurred during this time. However the mechanism underlying this difference in wild type MICs remains unknown. At present, the co-occurrence of resistance to both cephalosporins and fluoroquinolones hinders attempts to understand their relative importance in epidemic spread. Further controlled studies of cephalosporin and fluoroquinolone stewardship are needed.

Given the large number of PBP substituted, clinically important MDR clades which have emerged in the last 30 years, in different geographic regions (Figures 4-8), it is surprising that their elevated cephalosporin MICs have not been highlighted previously. This likely reflects the ‘intrinsic cephalosporin resistance’ concept (10), established in the early 1980s, when measured *C. difficile* cephalosporin MICs rarely exceeded 256µg/ml (33-36). These baseline MICs, which predate the emergence of the MDR clades, are already high relative to other pathogens, and were thought to exceed clinically relevant concentrations. The lack of a known mechanism of acquired β-lactam resistance compounded the situation.

Our phylogenies revealed the sequence and timing of MDR acquisition by clinically important lineages. The co-occurrence of PBP substitutions and fluoroquinolone resistance predated epidemic spread, which was reflected in short-branched, geographically structured clades (Figures 4-7). The first PBP substitution was typically PBP3 V497L, followed by others, yielding a variety of final combinations. The presence of *ermB*, (clindamycin resistance) and *rpoB* substitutions (rifampin resistance) was variable overall, but greater numbers of PBP substitutions frequently occurred with these in addition to fluoroquinolone resistance (Table S1, ST17(018, ST37(017)/ST81(369), ST8(002)/ST81).

We dated the emergence of the two MDR ST1(027) clades (FQ-R1 and FQ-R2) to the mid/late-1990s, as previously described (19) (Figure 4A,B). The emergence date of the MDR ST17(018) clade was compatible with first reports of outbreaks in 1996-1999 (59), the phylogeny root having a 95% credible interval dating of December 1998-July 2002 (Figure 5A). European and Asian ST17(018) clades then diverged, acquiring further region-specific PBP substitutions (Figure 5A), arguing against recent their recent intercontinental spread.

Greater numbers of PBP substitutions were associated with the highest cephalosporin MICs (Figure 3A). Consistent with this, highly substituted clades of multiple lineages (ST17(018), ST81(369)/ST37(018) and ST183/ST8(002, Figures 5A, 6A,B) predominated in South East Asia, where cephalosporin use is high (60-62). Adaptation to local prescribing conditions through PBP substitution acquisition offers a possible explanation for the temporal and geographic variation in prevalent *C. difficile* lineages (6, 8, 19, 47, 58, 63). This may extend to competitive exclusion of lesser PBP substituted strains by more highly substituted ones, a scenario requiring greater numbers of PBP substitutions to carry a fitness cost. This appears possible as in *C. perfringens in vitro*, PBP substitutions are associated with slower growth (64). We hypothesise that local levels of cephalosporin prescribing determine the prevalent *C. difficile* strain(s) in a given region. For example, the unusually low levels of ST1(027) seen in Asia, (65) may reflect competitive exclusion, under local prescribing conditions, by the more highly PBP substituted clades which predominate here. The relative geographic restriction of ST1(027)FQ-R1 (to USA, South Korea, Germany), compared to more globally distributed FQ-R2 may also reflect the different PBP substitutions of the two clades (Figure 4), and variations in MIC (Figure 3A).

MDR strains exhibit high transmissibility in clinical settings when prescribing is uncontrolled (17). The PBP1 and PBP3 transpeptidases function in cell wall biosynthesis, therefore substitutions impacting their catalytic domain could affect transmissibility via sporulation. A high sporulation phenotype has been reported in at least two epidemic lineages; ST3(001) (UK), and ST81(369) (Asia), (66, 67). Sporulation phenotype is reportedly variable in ST1(027), (68, 69), and we have observed variation in its PBP substitutions (Figure 4A,B, Table S1). However, the possibility of a link remains to be investigated. It is relevant to future experimental design that the MDR laboratory reference strains CD630 (ST54(012) and R20291 (ST1(027)) both contain PBP substitutions (Table 1).

PBP substitutions occurred *without* fluoroquinolone resistance in reasonable numbers of genomes of only three toxigenic lineages studied; ST2(020/014), ST54(012), and ST42(106) (USA) (Table S1). The latter was of interest since it recently exceeded the prevalence of ST1(027) in North America (8, 71). Visual inspection of phylogenies was used to assess whether PBP substitutions might enhance transmissibility in the absence of fluoroquinolone resistance. An ST2(020/014) phylogeny showed some possibility of locally enhanced transmission (Figure 8), as did ST54(012), and ST42(106) (USA) (Figure 7A,B). However, these events were small scale and this question remains to be answered. PBP substitutions (without fluoroquinolone resistance) were, however, widespread within the two non-toxigenic lineages ST15(010) and ST26(039), (Table S2) potentially explaining their high prevalence over other non-toxigenic strains.

The cephalosporin MICs for PBP substituted strains were extremely high for certain antibiotics (for example >512µg/ml for cefotaxime, and up to 1506µg/ml for cefuroxime, Figure 3A). This raises questions about the *in vivo* conditions required for PBP substitution selection. Intravenous β-lactams are eliminated in active form by biliary excretion, resulting in highly variable intestinal concentrations (71). Intestinal concentrations ranging from 1.01 to 1,345μg/ml have been reported (72) and so the potential exists for *C. difficile* to be exposed *in vivo* to cephalosporin concentrations reaching the MICs measured here. Resistant bacteria can also be selected experimentally at antimicrobial concentrations up to several hundred-fold below lethal levels (73-77). However, the overall contribution made by such ‘sub-MIC selection’ to resistance in clinically important bacteria is unknown (78).

To date, only ∼1% of known *C. difficile* lineages (949 STs identified as at 13 May 2022, https://pubmlst.org/organisms/clostridioides-difficile) have evolved MDR clade(s).

Furthermore, members of this minority have tended to evolve >1 such clade (Figure 4-7). This suggests a wild-type phenotype which favours the acquisition of chromosomal SNPs which raise cephalosporin and fluoroquinolone MICs. As discussed above, the baseline cephalosporin MICs for MDR-yielding lineages were higher than those lacking MDR strains (Figure 3A). This potentially favours survival of MDR-yielding lineages in low cephalosporin concentrations, allowing selection of PBP substitutions. An alternative, mechanism might be a hypermutator phenotype, as in *S. pneumoniae* (73).

In summary, our findings identify a role for cephalosporin selection in the evolution of epidemic CDI lineages. Specific regional prescribing practises may determine the locally predominant epidemic strains, potentially explaining the marked international variation in *C. difficile* molecular epidemiology. Since antimicrobial stewardship typically targets multiple drug classes (29, 79) and epidemic strains have raised MICs for fluoroquinolones, cephalosporins, (Figure 2) and more variably clindamycin (*ermB*) (Table S1), is difficult to determine the relative contributions made by stewardship of each drug to CDI control. The timing of cephalosporin and fluoroquinolone resistance acquisition, immediately before the emergence of multiple epidemic strains from divergent *C. difficile* genetic backgrounds, suggests AMR may be equally important as, or even exceed, strain-specific virulence determinants in driving epidemic CDI.

## MATERIALS AND METHODS

WGS from 7094 *C. difficile* isolates, predominantly cultured from humans with CDI were obtained. Clinical isolates from hospital and community patients from Europe, North and South America, South East Asia, and Australia were included. Fourteen CDI lineages were represented, and two non-toxigenic lineages. A complete list of genomes, their identifiers in public databases, and references is provided (Table S1). Raw sequence reads were assembled *de novo* as required using Velvet (version 1.0.7 – 1.0.18) (80) and Velvet Optimiser with default settings (2.1.7) (81). A minority of genomes were obtained assembled, either from the NCBI database (82) or EnteroBase (83, 84). Assemblies were imported to a BIGSdb database (85) which was used to identify the seven loci used in multi-locus sequence typing (44). Sequence types (STs) were assigned using the *C. difficile* PubMLST database (https://pubmlst.org/organisms/clostridioides-difficile/). ST and PCR-ribotype were used to indicate genetic lineages, identified by the notation ST1(027) (sequence type-1 (PCR-ribotype-027)).

BLAST searches performed within BIGSdb (85) were used to identify and extract chromosomal gene sequences for PBP transpeptidases (PBP1-5, (45)), together with *gyrA*, *gyrB*, and *rpoB*, specific mutations in which confer AMR. Established amino acid substitutions scored as conferring resistance to fluoroquinolones were GyrA T81I and GyrB D426N, and to rifampin RpoB R505K, H502N, S498T. Acquisition of *ermB*, conferring clindamycin resistance was also noted (86-89). Each unique allele sequence identified at these loci (PBP1-5, *gyrA*, *gyrB*, *rpoB* and *ermB*) was assigned a number (Table S1) and can be downloaded at https://pubmlst.org/organisms/clostridioides-difficile/ (44, 85). Newly extracted gene sequences were queried against this database and the allele numbers were recorded for each genome, together with the substitutions relevant to AMR (Table S1).

### Identification of recent PBP substitutions

This was achieved using MEGA (https://www.megasoftware.net/) (90), which facilitated within-lineage comparisons of the nucleotide and amino acid sequences of PBP1-5 alleles. Comparisons were made relative to the wild type PBP sequence for each lineage, wild-type alleles being taken from non-MDR genomes within the lineage.

### Phenotyping

Isolates were chosen for phenotyping from four lineages, each of which contained both PBP substituted and wild type strains; ST1(027)FQ-R1 and FQ-R2, ST3(001), ST17(017) and ST42(106). Representatives for each lineage were chosen on the basis of minimal SNP distances (Figure 3A). Genomes of four lineages lacking MDR strains (wild type only), also underwent phenotyping; ST10(015), ST6(005), ST56(058) and ST7(026).

Minimum inhibitory concentrations of cefotaxime, cefuroxime, cephradine, amoxicillin, amoxicillin-clavulante, meropenem, imipenem and piperacillin-tazobatam were determined by Wilkins Chalgren agar dilution methods (91, 92). Briefly, *C. difficile* isolates and controls (*C. difficile* ATCC700057, E4 (PCR ribotype 010) and *B. fragilis* 25285) were cultured in pre-reduced Schaedlers anaerobic broths at 37°C for 24h, anaerobically. Isolates and controls were diluted in pre-reduced saline to McFarland standard 1 equivalence and multipoint inoculated onto prepared antibiotic-containing agar plates and controls. Agar plates were incubated at 37°C for 24h, anaerobically prior to MIC determination. MIC was defined as the lowest concentration of antimicrobial that completely inhibited growth, showed only 1 or 2 colonies, or left a faint haze of growth on the plate.

Antimicrobial concentrations were prepared using solvents and diluents recommended in the CLSI guidelines (93, 94). For amoxicillin clavulanate and piperacillin-tazobactam, clavulanic acid and tazobactam were was added to agar at fixed concentrations of 2mg/L and 4 mg/L respectively. In order to test susceptibility within normal doubling dilutions, further antibiotic concentrations were prepared for the antibiotic plate range. All antibiotics were tested at the following dilutions: 0.125, 0.25, 0.5, 1, 2, 4, 8, 16, 32, 36, 40, 46, 53, 64, 71, 80, 91, 107, 128, 142, 160, 182, 213, 256, 284, 320, 366, 427, 512 mg/L. Additionally, cefuroxime and cephradine were prepared up to their limit of solubility and the following ranges of dilutions were prepared. Cefuroxime: 102, 120, 143, 160, 205, 213, 240, 287, 319, 409, 478, 572 mg/L. Cephradine: 105, 118, 134, 157, 188, 209, 235, 269, 314, 376, 418, 471, 538, 627, 753, 837, 941, 1076, 1255, 1506 mg/L.

### Construction of dated phylogenies

Dated phylogenies were constructed using genomes chosen to maximise geographic and temporal spread of wild type and AMR strains, as well as to represent the diversity of PBP substitutions detected in each lineage. Each set of genomes was first aligned to a reference using MuMMER version 3.1 (95) to produce a genome-wide alignment. Initial phylogenies were built using PhyML version 3.3 (96) which were then corrected for recombination using ClonalFrameML version 1.12 (97). Finally these phylogenies were dated using BactDating version 1.1 (98) assuming a mean evolutionary rate of 1.4 mutations per year per genome as in previous similar studies (99).

## ACKNOWLEDGEMENTS

This study was supported by the National Institute for Health Research (NIHR) Oxford Biomedical Research Centre (BRC), and by the National Institute for Health Research (NIHR) Health Protection Research Unit in Healthcare Associated Infections and Antimicrobial Resistance (NIHR200915), a partnership between the UK Health Security Agency (UKHSA) and the University of Oxford.

XD was funded by the NIHR Health Protection Research Unit (HPRU) in Genomics and Enabling Data, and the NIHR HPRU in Gastrointestinal Infections.

MHW is supported by the National Institute for Health Research Leeds in Vitro Diagnostics Co-operative.

DWC is a NIHR Senior Investigator ASW is a NIHR Senior Investigator. DWE is a Robertson Foundation Fellow.

The funders had no role in study design, data collection and interpretation, or the decision to submit the work for publication. The views expressed are those of the authors and not necessarily those of the NHS, NIHR or Department of Health.

## DECLARATIONS

DWE declares lecture fees from Gilead, outside the submitted work. No other author has a conflict of interest to declare.

## SUPPLEMENTAL MATERIAL

**Figure S1.** ST81(369) PBP3 was acquired via a long recombination event.

Distance plots showing pairwise comparisons of the donor ST8(002), recipient ST37(017) and recombinant ST81(369), with the aim of identifying the size and location of the recombination event. The genomes used were ST8(002) isolate W0003a NZ_CP025047.1 (Yin et al., 2018), ST37(017) isolate M68 NC_017175.1 (He et al., 2010), and ST81(369) isolate 28 WGS:QNWI01, (Bioproject PRJNA479396, Assembly GCA_003326885.1) (Wu et al., 2019).

The plots extend from 0.9 to 1.4 Mbp relative to M68 (x axis) so the locations of both PBP1 and PBP3 are covered; PBP1 at 907,056-904,363 (gene CDM68_RS04280) and PBP3 at 1,219,021-1,221,999, (gene CDM68_RS05670), indicated by vertical red dashed lines.

Missing lines indicate alignment gaps.

The large recombination event involves ∼150kbp (∼3.5% of the 4,308,325bp genome). A much smaller region occurred near the middle of the large event (position ∼1.22Mbp) where ST81(369) no longer resembles ST8(002), but resembles ST37(017) instead. It appears likely that this short region, adjacent to PBP3, has recombined back with ST37(017).

## REFERENCES

1. Smits WK, Lyras D, Lacy DB, Wilcox MH, Kuijper EJ. 2016. *Clostridium difficile* infection. Nat Rev Dis Primers 2:16020.

2. McDonald LC, Killgore GE, Thompson A, Owens RC Jr, Kazakova SV, Sambol SP, Johnson S, Gerding DN. 2005. An epidemic, toxin gene-variant strain of *Clostridium difficile*. N Engl J Med 353:2433–2441.

3. Zaiss NH, Witte, W, Nübel U. 2010. Fluoroquinolone resistance and *Clostridium difficile*, Germany. Emerg Infect Dis 16:675–677.

4. Barbanti F, Spigaglia P. 2016. Characterization of *Clostridium difficile* PCR-ribotype 018: A problematic emerging type. Anaerobe 42:123–129.

5. Imwattana K, Knight DR, Kullin B, Collins DA, Putsathit P, Kiratisin P, Riley TV. 2020. Antimicrobial resistance in *Clostridium difficile* ribotype 017. Expert Rev Anti Infect Ther 18:17–25.

6. Freeman J, Vernon J, Pilling S, Morris K, Nicolson S, Shearman S, Clark E, Palacios-Fabrega JA, Wilcox M; Pan-European Longitudinal Surveillance of Antibiotic Resistance among Prevalent *Clostridium difficile* Ribotypes’ Study Group. 2020. Five-year Pan-European, longitudinal surveillance of *Clostridium difficile* ribotype prevalence and antimicrobial resistance: the extended ClosER study. Eur J Clin Microbiol Infect Dis 39:169-177.

7. Lew T, Putsathit P, Sohn KM, Wu Y, Ouchi K, Ishii Y, Tateda K, Riley TV, Collins DA. 2020. Antimicrobial Susceptibilities of *Clostridium difficile* Isolates from 12 Asia-Pacific Countries in 2014 and 2015. Antimicrob Agents Chemother 64:e00296–20.

8. Carlson TJ, Blasingame D, Gonzales-Luna AJ, Alnezary F, Garey KW. 2020. *Clostridioides difficile* ribotype 106: A systematic review of the antimicrobial susceptibility, genetics, and clinical outcomes of this common worldwide strain. Anaerobe 62:102142.

9. Owens RC Jr, Donskey CJ, Gaynes RP, Loo VG, Muto CA. 2008. Antimicrobial-associated risk factors for *Clostridium difficile* infection. Clin Infect Dis 46 Suppl 1:S19–31.

10. Gerding DN. 2004. Clindamycin, cephalosporins, fluoroquinolones, and *Clostridium difficile*-associated diarrhea: this is an antimicrobial resistance problem. Clin Infect Dis 38:646–648.

11. Muto CA, Pokrywka M, Shutt K, Mendelsohn AB, Nouri K, Posey K, Roberts T, Croyle K, Krystofiak S, Patel-Brown S, Pasculle AW, Paterson DL, Saul M, Harrison LH. 2005. A large outbreak of *Clostridium difficile*-associated disease with an unexpected proportion of deaths and colectomies at a teaching hospital following increased fluoroquinolone use. Infect Control Hosp Epidemiol 26:273–280.

12. Johnson S, Samore MH, Farrow KA. 1999. Epidemics of diarrhea caused by a clindamycin-resistant strain of *Clostridium difficile* in four hospitals. N Engl J Med 341:1645–1651.

13. Loo VG, Poirier L, Miller MA, Oughton M, Libman MD, Michaud S, Bourgault AM, Nguyen T, Frenette C, Kelly M, Vibien A, Brassard P, Fenn S, Dewar K, Hudson TJ, Horn R, René P, Monczak Y, Dascal A. 2005. A predominantly clonal multi-institutional outbreak of *Clostridium difficile*-associated diarrhea with high morbidity and mortality. N Engl J Med 353:2442–2449.

14. Arvand M, Hauri AM, Zaiss NH, Witte W, Bettge-Weller G. 2009. *Clostridium difficile* ribotypes 001, 017, and 027 are associated with lethal *C. difficile* infection in Hesse, Germany. Euro Surveill 14:19403.

15. Goorhuis A, DebastSB, Dutilh JC, van Kinschot CM, Harmanus C, Cannegieter SC, Hagen EC, Kuijper EJ. 2011. Type-specific risk factors and outcome in an outbreak with two different *Clostridium difficile* types simultaneously in one Hospital. Clinical Infectious Diseases 53:860–869.

16. Walker AS, Eyre DW, Wyllie DH, Dingle KE, Griffiths D, Shine B, Oakley S, O’Connor L, Finney J, Vaughan A, Crook DW, Wilcox MH, Peto TE; Infections in Oxfordshire Research Database. 2013. Relationship between bacterial strain type, host biomarkers, and mortality in *Clostridium difficile* infection Clin Infect Dis 56:1589-1600.

17. Baldan R, Trovato A, Bianchini V, Biancardi A, Cichero P, Mazzotti M, Nizzero P, Moro M, Ossi C, Scarpellini P, Cirillo DM. 2015. *Clostridium difficile* PCR Ribotype 018, a Successful Epidemic Genotype J Clin Microbiol 53:2575-2580.

18. Serafino S, Consonni D, Migone De Amicis M, Sisto F, Domeniconi G, Formica S, Zarantonello M, Maraschini A, Cappellini MD, Spigaglia P, Barbanti F, Castaldi S, Fabio G. 2018. Clinical outcomes of *Clostridium difficile* infection according to strain type. A prospective study in medical wards. Eur J Intern Med 54:21–26.

19. He M, Miyajima F, Roberts P, Ellison L, Pickard DJ, Martin MJ, Connor TR, Harris SR, Fairley D, Bamford KB, D’Arc S, Brazier J, Brown D, Coia JE, Douce G, Gerding D, Kim HJ, Koh TH, Kato H, Senoh M, Louie T, Michell S, Butt E, Peacock SJ, Brown NM, Riley T, Songer G, Wilcox M, Pirmohamed M, Kuijper E, Hawkey P, Wren BW, Dougan G, Parkhill J, Lawley TD. 2013. Emergence and global spread of epidemic healthcare-associated *Clostridium difficile*. Nat Genet 45:109–113.

20. Senoh M, Haru Kato H. 2022. Molecular epidemiology of endemic *Clostridioides difficile* infection in Japan. Anaerobe 3:102510.

21. Hensgens MP, Goorhuis A, van Kinschot CM, Crobach MJ, Harmanus C, Kuijper EJ. 2011. *Clostridium difficile* infection in an endemic setting in the Netherlands. Eur J Clin Microbiol Infect Dis 30:587–93.

22. Eyre DW, Davies KA, Davis G, Fawley WN, Dingle KE, De Maio N, Karas A, Crook DW, Peto TEA, Walker AS, Wilcox MH; EUCLID Study Group. 2018. Two distinct patterns of *Clostridium difficile* diversity across Europe indicating contrasting routes of spread. Clin Infect Dis 67:1035-1044.

23. Pear SM, Williamson TH, Bettin KM, Gerding DN, Galgiani JN. 1994. Decrease in nosocomial *Clostridium difficile*-associated diarrhea by restricting clindamycin use. Ann Intern Med 120:272–277.

24. McNulty C, Logan M, Donald IP, Ennis D, Taylor D, Baldwin RN, Bannerjee M, Cartwright KA. 1997. Successful control of *Clostridium difficile* infection in an elderly care unit through use of a restrictive antibiotic policy J Antimicrob Chemother 40:707–711.

25. Valiquette L, Cossette B, Garant MP, Diab H, Pépin J. 2007. Impact of a reduction in the use of high-risk antibiotics on the course of an epidemic of *Clostridium difficile*-associated disease caused by the hypervirulent NAP1/027 strain. Clin Infect Dis 45:S112–S121.

26. Debast SB, Vaessen N, Choudry A, Wiegers-Ligtvoet EAJ, van den Berg RJ, Kuijper EJ. 2009. Successful combat of an outbreak due to *Clostridium difficile* PCR ribotype 027 and recognition of specific risk factors Clin Microbiol Infect 15:427-434.

27. Feazel LM, Malhotra A, Perencevich EN, Kaboli P, Diekema DJ, Schweizer ML. 2014. Effect of antibiotic stewardship programmes on *Clostridium difficile* incidence: a systematic review and meta-analysis. J Antimicrob Chemother 69:1748–1754.

28. Sarma JB, Marshall B, Cleeve V, Tate D, Oswald T, Woolfrey S. 2015. Effects of fluoroquinolone restriction (from 2007 to 2012) on *Clostridium difficile* infections: interrupted time-series analysis. J Hosp Infect 91:74–80.

29. Muto CA, Blank MK, Marsh JW. 2007. Control of an outbreak of infection with the hypervirulent *Clostridium difficile* BI strain in a university hospital using a comprehensive “bundle” approach. Clin Infect Dis 45:1266–1273.

30. Dingle KE, Didelot X, Quan TP, Eyre DW, Stoesser N, Golubchik T, Harding RM, Wilson DJ, Griffiths D, Vaughan A, Finney JM, Wyllie DH, Oakley SJ, Fawley WN, Freeman J, Morris K, Martin J, Howard P, Gorbach S, Goldstein EJC, Citron DM, Hopkins S, Hope R, Johnson AP, Wilcox MH, Peto TEA, Walker AS, Crook DW; Modernising Medical Microbiology Informatics Group. 2017. Effects of control interventions on *Clostridium difficile* infection in England: an observational study. Lancet Infect Dis 17:411-421.

31. Lawes T, Lopez-Lozano JM, Nebot CA, Macartney G, Subbarao-Sharma R, Wares KD, Sinclair C, Gould IM. 2017. Effect of a national 4C antibiotic stewardship intervention on the clinical and molecular epidemiology of *Clostridium difficile* infections in a region of Scotland: a non-linear time-series analysis. Lancet Infect Dis 17:194–206.

32. Huang H, Weintraub A, Fang H, Nord CE. 2009. Antimicrobial resistance in *Clostridium difficile*. Int J Antimicrob Agents 34:516–522.

33. Dzink J, Bartlett JG. 1980. *In vitro* susceptibility of *Clostridium difficile* isolates from patients with antibiotic-associated diarrhea or colitis. Antimicrob Agents Chemother 17:695–698.

34. Shuttleworth R, Taylor M, Jones DM. 1980. Antimicrobial susceptibilities of *Clostridium difficile*. J Clin Pathol 33:1002–1005.

35. Greenfield RA, Kurzynski TA, Craig, WA. 1982. *In vitro* susceptibility of *Clostridium difficile* isolates to cefotaxime, moxalactam, and cefoperazone. Antimicrob Agents and Chemother 21:846–847.

36. Chow AW, Cheng N, Bartlett KH. 1985. *In vitro* susceptibility of *Clostridium difficile* to new beta-lactam and quinolone antibiotics. Antimicrob Agents Chemother 28:842–844.

37. Toth M, Stewart NK, Smith C, Vakulenko SB. 2018. Intrinsic Class D β-Lactamases of *Clostridium difficile*. mBio 9:e01803–18.

38. Sandhu BK, Edwards AN, Anderson SE, Woods EC, McBride SM. 2019. Regulation and Anaerobic Function of the *Clostridioides difficile* β-Lactamase. Antimicrob Agents Chemother 64:e01496–19.

39. Fisher JF, Mobashery S. 2016. β-Lactam Resistance Mechanisms: Gram-Positive Bacteria and *Mycobacterium tuberculosis*. Cold Spring Harb Perspect Med 6:a025221.

40. Sauvage E, Kerff F, Terrak M, Ayala JA, Charlier P. 2008. The penicillin-binding proteins: structure and role in peptidoglycan biosynthesis. FEMS Microbiology Reviews 32:234–258.

41. Zapun A, Contreras-Martel C, Vernet T. 2008. Penicillin-binding proteins and β-lactam resistance. FEMS Microbiol Rev 32:361–385.

42. Kwon DH, Dore MP, Kim JJ, Kato M, Lee M, Wu JY, Graham DY. 2003. High-level beta-lactam resistance associated with acquired multidrug resistance in *Helicobacter pylori*. Antimicrob Agents Chemother 47:2169–2178.

43. Dewé TCM, D’Aeth JC, Croucher NJ. 2019. Genomic epidemiology of penicillin-non-susceptible *Streptococcus pneumoniae*. Microb Genom 5:e000305.

44. Griffiths D, Fawley W, Kachrimanidou M, Bowden R, Crook DW, Fung R, Golubchik T, Harding RM, Jeffery KJ, Jolley KA, Kirton R, Peto TE, Rees G, Stoesser N, Vaughan A, Walker AS, Young BC, Wilcox M, Dingle KE. 2010. Multilocus sequence typing of *Clostridium difficile*. J Clin Microbiol 483:770–778.

45. Isidro J, Santos A, Nunes A, Borges V, Silva C, Vieira L, Mendes AL, Serrano M, Henriques AO, Gomes JP, Oleastro M. 2018. Imipenem Resistance in *Clostridium difficile* Ribotype 017, Portugal. Emerg Infect Dis 24:741–745.

46. Dembek M, Barquist L, Boinett CJ, Cain AK, Mayho M, Lawley TD, Fairweather NF, Fagan RP. 2015. High-throughput analysis of gene essentiality and sporulation in *Clostridium difficile*. mBio 6:e02383.

47. Aoki K, Takeda S, Miki T, Ishii Y, Tateda K. 2019. Antimicrobial susceptibility and molecular characterisation using whole-genome sequencing of *Clostridioides difficile* collected in 82 hospitals in Japan between 2014 and 2016. Antimicrob Agents Chemother. 63:e01259–19.

48. Jia H, Du P, Yang H, Zhang Y, Wang J, Zhang W, Han G, Han N, Yao Z, Wang H, Zhang J, Wang Z, Ding Q, Qiang Y, Barbut F, Gao GF, Cao Y, Cheng Y, Chen C. 2016. Nosocomial transmission of *Clostridium difficile* ribotype 027 in a Chinese hospital, 2012-2014, traced by whole genome sequencing. BMC Genomics 17:405.

49. Qin J, Dai Y, Ma X, Wang Y, Gao Q, Lu H, Li T, Meng H, Liu Q, Li M. 2017. Nosocomial transmission of *Clostridium difficile* Genotype ST81 in a general teaching hospital in China traced by whole genome sequencing. Sci Rep 7:9627.

50. Wu Y, Liu C, Li WG, Xu JL, Zhang WZ, Dai YF, Lu JX. 2019. Independent Microevolution Mediated by Mobile Genetic Elements of Individual *Clostridium difficile* Isolates from Clade 4 Revealed by Whole-Genome Sequencing. mSystems 4:e00252–18.

51. Ramírez-Vargas G, Quesada-Gómez C, Acuña-Amador L, López-Ureña D, Murillo T, Del Mar Gamboa-Coronado M, Chaves-Olarte E, Thomson N, Rodríguez-Cavallini E, Rodríguez C. 2017. A *Clostridium difficile* lineage endemic to Costa Rican hospitals is multidrug resistant by acquisition of chromosomal mutations and novel mobile genetic elements. Antimicrob Agents Chemother 61:e02054-16.

52. Kachrimanidou M, Baktash A, Metallidis S, Tsachouridou Ο, Netsika F, Dimoglou D, Kassomenaki A, Mouza E, Haritonidou M, Kuijper E. 2020. An outbreak of *Clostridioides difficile* infections due to a 027-like PCR ribotype 181 in a rehabilitation centre: Epidemiological and microbiological characteristics. Anaerobe 65:102252.

53. Boekhoud IM, Sidorov I, Nooij S, Harmanus C, Bos-Sanders IMJG, Viprey V, Spittal W, Clark E, Davies K, Freeman J, Kuijper EJ, Smits WK; COMBACTE-CDI Consortium. 2021. Haem is crucial for medium-dependent metronidazole resistance in clinical isolates of *Clostridioides difficile*. J Antimicrob Chemother 76:1731-1740.

54. Imwattana K, Putsathit P, Knight DR, Kiratisin P, Riley TV. 2021. Molecular Characterization of, and Antimicrobial Resistance in, *Clostridioides difficile* from Thailand, 2017-2018. Microb Drug Resist 27:1505-1512.

55. Hiramatsu K, Cui L, Kuroda M, Ito T. 2001. The emergence and evolution of methicillin-resistant *Staphylococcus aureus*. Trends Microbiol 9:486–493.

56. Ludlam H, Brown N, Sule O, Redpath C, Coni N, Owen G. 1999. An antibiotic policy associated with reduced risk of *Clostridium difficile*-associated diarrhoea. Age Ageing 28:578–580.

57. O’Connor KA, Kingston M, O’Donovan M, Cryan B, Twomey C, O’Mahony D. 2004. Antibiotic prescribing policy and *Clostridium difficile* diarrhoea QJM. 97:423–429.

58. Belmares J, Johnson S, Parada JP. 2009. Molecular epidemiology of *Clostridium difficile* over the course of 10 years in a tertiary care hospital. Clin Infect Dis 49:1141–1147.

59. Kato H, Kita H, Karasawa T, Maegawa T, Koino Y, Takakuwa H, Saikai T, Kobayashi K, Yamagishi T, Nakamura S. 2001. Colonisation and transmission of *Clostridium difficile* in healthy individuals examined by PCR ribotyping and pulsed-field gel electrophoresis. J Med Microbiol 50:720–727.

60. Muraki Y, Yagi T, Tsuji Y, Nishimura N, Tanabe M, Niwa T, Watanabe T, Fujimoto S, Takayama K, Murakami N, Okuda M. 2016. Japanese antimicrobial consumption surveillance: First report on oral and parenteral antimicrobial consumption in Japan (2009-2013). J Glob Antimicrob Resist 7:19–23.

61. Kim YA, Park YS, Youk T, Lee H, Lee K. 2018. Changes in Antimicrobial Usage Patterns in Korea: 12-Year Analysis Based on Database of the National Health Insurance Service-National Sample Cohort. Sci Rep 81:12210.

62. Qu X, Yin C, Sun X, Huang S, Li C, Dong P, Lu X, Zhang Z, Yin A. 2018. Consumption of antibiotics in Chinese public general tertiary hospitals (2011-2014): Trends, pattern changes and regional differences. PLoS One 13:e0196668.

63. Collins DA, Hawkey PM, Riley TV. 2013. Epidemiology of *Clostridium difficile* infection in Asia. Antimicrob Resist Infect Control 2:21.

64. Park M, Rafii F. 2017. Exposure to beta-lactams results in the alteration of penicillin-binding proteins in *Clostridium perfringens*. Anaerobe 45:78–85.

65. Cheng JW, Xiao M, Kudinha T, Xu Z, Hou X, Sun L, Zhang L, Fan X, Kong F, Xu Y. 2016. The first two *Clostridium difficile* Ribotype 027/ST1 Isolates identified in Beijing, China-an emerging problem or a neglected threat? Sci Rep 6:18834.

66. Wilcox MH, Fawley WN. 2000. Hospital disinfectants and spore formation by *Clostridium difficile*. Lancet 356:1324.

67. Wang B, Peng W, Zhang P, Su J. 2018. The characteristics of *Clostridium difficile* ST81, a new PCR ribotype of toxin A-B+ strain with high-level fluoroquinolones resistance and higher sporulation ability than ST37/PCR ribotype 017. FEMS Microbiol Lett 365(17).

68. Burns DA, Heap JT, Minton NP. 2010. The diverse sporulation characteristics of *Clostridium difficile* clinical isolates are not associated with type. Anaerobe 16:618–622.

69. Burns DA, Heeg D, Cartman ST, Minton NP. 2011. Reconsidering the sporulation characteristics of hypervirulent *Clostridium difficile* BI/NAP1/027. PLoS One 6:e24894.

70. Karlowsky JA, Adam HJ, Baxter MR, Dutka CW, Nichol KA, Laing NM, Golding GR, Zhanel GG. 2020. Antimicrobial susceptibility of *Clostridioides difficile* isolated from diarrhoeal stool specimens of Canadian patients: summary of results from the Canadian *Clostridioides difficile* (CAN-DIFF) surveillance study from 2013 to 2017. J Antimicrob Chemother 75:1824–1832.

71. Karachalios G, Charalabopoulos K. 2002. Biliary excretion of antimicrobial drugs. Chemotherapy 48:280–297.

72. Kokai-Kun JF, Roberts T, Coughlin O, Sicard E, Rufiange M, Fedorak R, Carter C, Adams MH, Longstreth J, Whalen H, Sliman J. 2017. The oral beta-lactamase SYN-004 (ribaxamase) degrades ceftriaxone excreted into the intestine in phase 2a clinical studies. Antimicrob Agents Chemother 61:e02197–16.

73. Negri MC, Morosini MI, Baquero MR, del Campo R, Blázquez J, Baquero F. 2002. Very low cefotaxime concentrations select for hypermutable *Streptococcus pneumoniae* populations. Antimicrob Agents Chemother 46:528-530.

74. Gullberg E, Cao S, Berg OG, Ilbäck C, Sandegren L, Hughes D, Andersson DI. 2011. Selection of resistant bacteria at very low antibiotic concentrations. PLoS Pathog 7:e1002158.

75. Gullberg E, Albrecht LM, Karlsson C, Sandegren L, Andersson DI. 2014. Selection of a multidrug resistance plasmid by sublethal levels of antibiotics and heavy metals. mBio 5: e01918–01914.

76. Andersson DI, Hughes D. 2012. Evolution of antibiotic resistance at non-lethal drug concentrations. Drug Resist Updat 15:162–172.

77. Murray AK, Zhang L, Yin X, Zhang T, Buckling A, Snape J, Gaze WH. 2018. Novel insights into selection for antibiotic resistance in complex microbial communities. mBio 9:e00969–18.

78. Sandegren L. 2014. Selection of antibiotic resistance at very low antibiotic concentrations. Ups J Med Sci 119:103–107.

79. http://www.hpa.org.uk/webc/HPAwebFile/HPAweb_C/1232006607827

80. Zerbino DR, Birney E. 2008. Velvet: algorithms for de-novo short read assembly using de Bruijn graphs. Genome Res 18:821–829.

81. Gladman S, Seemann T. 2008. VelvetOptimiser, 2.1.7. Monash University, Victoria, Australia.

82. NCBI database. https://www.ncbi.nlm.nih.gov/genome/browse#!/prokaryotes/535/

83. Enterobase https://enterobase.warwick.ac.uk/species/index/clostridium

84. Frentrup M, Zhou Z, Steglich M, Meier-Kolthoff JP, Göker M, Riedel T, Bunk B, Spröer C, Overmann J, Blaschitz M, Indra A, von Müller L, Kohl TA, Niemann S, Seyboldt C, Klawonn F, Kumar N, Lawley TD, García-Fernández S, Cantón R, Del Campo R, Zimmermann O, Groß U, Achtman M, Nübel U. 2020. A publicly accessible database for *Clostridioides difficile* genome sequences supports tracing of transmission chains and epidemics. Microb Genom 6:mgen000410.

85. Jolley KA, Bray JE, Maiden MCJ. 2018. Open-access bacterial population genomics: BIGSdb software, the PubMLST.org website and their applications. Wellcome Open Res 3:124.

86. Spigaglia P, Barbanti F, Mastrantonio P. 2008. Fluoroquinolone resistance in *Clostridium difficile* isolates from a prospective study of *C. difficile* infections in Europe. J Med Microbiol 57:784–789.

87. Drudy D, Quinn T, O’Mahony R, Kyne L, O’Gaora P, Fanning S. 2006. High-level resistance to moxifloxacin and gatifloxacin associated with a novel mutation in *gyrB* in toxin-A-negative, toxin-B-positive *Clostridium difficile*. J Antimicrob Chemother 58:1264-1267.

88. Curry SR, Marsh JW, Shutt KA, Muto CA, O’Leary MM, Saul MI, Pasculle AW, Harrison LH. 2009. High frequency of rifampin resistance identified in an epidemic *Clostridium difficile* clone from a large teaching hospital. Clin Infect Dis 48:425–429.

89. Spigaglia P, Barbanti F, Mastrantonio P; European Study Group on Clostridium difficile (ESGCD). 2011. Multidrug resistance in European *Clostridium difficile* clinical isolates. J Antimicrob Chemother 66:2227-2234.

90. Tamura K, Stecher G, Kumar S. 2021. MEGA11: Molecular Evolutionary Genetics Analysis Version 11. Molecular Biology and Evolution 38:3022–3027.

91. Baines SD, O’Connor R, Freeman J, Fawley WN, Harmanus C, Mastrantonio, Kuijper EJ, Wilcox MJ. 2008. Emergence of reduced susceptibility to metronidazole in *Clostridium difficile*. J Antimicrob Chemother 62:1046–1052.

92. Freeman, J, Vernon, J, Morris K, Nicholson S, Todhunter S, Longshaw C, Wilcox MH. 2015. Pan-European longitudinal surveillance of antibiotic resistance among prevalent *Clostridium difficile* ribotypes. Clin Microbiol Infect 21:e9–16.

93. Clinical Laboratory Standards Institute. M11-A8. Methods for Antimicrobial Susceptibility Testing of Anaerobic Bacteria; Approved Standard - 8th Edition. 2012.

94. Clinical Laboratory Standards Institute, M100S. Performance Standards for Antimicrobial Susceptibility Testing. 2012.

95. Kurtz S, Phillippy A, Delcher AL, Smoot M, Shumway M, Antonescu C, Salzberg SL. 2004. Versatile and open software for comparing large genomes. Genome Biol. 5:R12.

96. Guindon S, Dufayard J-F, Lefort V, Anisimova M, Hordijk W, Gascuel O. 2010. New algorithms and methods to estimate maximum-likelihood phylogenies: assessing the performance of PhyML 3.0. Syst Biol 59:307–321.

97. Didelot X, Wilson DJ. 2015. ClonalFrameML: Efficient Inference of Recombination in Whole Bacterial Genomes. PLoS Comput Biol 11:e1004041.

98. Didelot X, Croucher NJ, Bentley SD, Harris SR, Wilson DJ. 2018. Bayesian inference of ancestral dates on bacterial phylogenetic trees. Nucleic Acids Res. 46:e134.

99. Didelot X, Eyre DW, Cule M, Ip CLC, Ansari MA, Griffiths D, Vaughan A, O’Connor L, Golubchik T, Batty EM, Piazza P, Wilson DJ, Bowden R, Donnelly PJ, Dingle KE, Wilcox M, Walker AS, Crook DW, Peto TE, Harding RM. 2012. Microevolutionary analysis of *Clostridium difficile* genomes to investigate transmission. Genome Biology 13:R118.

100. Lawley TD, Croucher NJ, Yu L, Clare S, Sebaihia M, Goulding D, Pickard DJ, Parkhill J, Choudhary J, Dougan G. 2009. Proteomic and genomic characterization of highly infectious *Clostridium difficile* 630 spores. J Bacteriol 191:5377–5386.

101. Pettit LJ, Browne HP, Yu L, Smits WK, Fagan RP, Barquist L, Martin MJ, Goulding D, Duncan SH, Flint HJ, Dougan G, Choudhary JS, Lawley TD. 2014. Functional genomics reveals that *Clostridium difficile* Spo0A coordinates sporulation, virulence and metabolism. BMC Genomics 15:160.

102. He M, Sebaihia M, Lawley TD, Stabler RA, Dawson LF, Martin MJ, Holt KE, Seth-Smith HM, Quail MA, Rance R, Brooks K, Churcher C, Harris D, Bentley SD, Burrows C, Clark L, Corton C, Murray V, Rose G, Thurston S, van Tonder A, Walker D, Wren BW, Dougan G, Parkhill J. 2010. Evolutionary dynamics of *Clostridium difficile* over short and long time scales. Proc Natl Acad Sci USA 107:7527–7532.

103. Sebaihia M, Wren BW, Mullany P, Fairweather NF, Minton N, Stabler R, Thomson NR, Roberts AP, Cerdeño-Tárraga AM, Wang H, Holden MT, Wright A, Churcher C, Quail MA, Baker S, Bason N, Brooks K, Chillingworth T, Cronin A, Davis P, Dowd L, Fraser A, Feltwell T, Hance Z, Holroyd S, Jagels K, Moule S, Mungall K, Price C, Rabbinowitsch E, Sharp S, Simmonds M, Stevens K, Unwin L, Whithead S, Dupuy B, Dougan G, Barrell B, Parkhill J. 2006. The multidrug-resistant human pathogen *Clostridium difficile* has a highly mobile, mosaic genome. Nat Genet 38:779–786.

104. Stoesser N, Eyre DW, Quan TP, Godwin H, Pill G, Mbuvi E, Vaughan A, Griffiths D, Martin J, Fawley W, Dingle KE, Oakley S, Wanelik K, Finney JM, Kachrimanidou M, Moore CE, Gorbach S, Riley TV, Crook DW, Peto TEA, Wilcox MH, Walker AS; Modernising Medical Microbiology Informatics Group (MMMIG). 2017. Epidemiology of *Clostridium difficile* in infants in Oxfordshire, UK: Risk factors for colonization and carriage, and genetic overlap with regional *C. difficile* infection strains. PLoS One. 12(8):e0182307.

